# A method for extracting an approximated connectome from libraries of single cell anatomical reconstructions

**DOI:** 10.1101/2023.05.21.541471

**Authors:** K.K.H. Manjunatha, M. Bruzzone, G. Nicoletti, S. Suweis, M. dal Maschio

**Affiliations:** Department of Biomedical Sciences, Università degli Studi di Padova, Padua, IT; Department of Physics and Astronomy, Università degli Studi di Padova, Padua, IT; Padova Neuroscience Center, Università degli Studi di Padova, Padua, IT; Modelling and Engineering Risk and Complexity, Scuola Superiore Meridionale, Napoli, IT

## Abstract

Functional brain activity is supported by specific circuit wiring diagrams. When the synaptic annotation is available, the analysis of high-resolution connectomes allows for unraveling the circuit architectures. Despite the continuous technological and computational improvements, obtaining a whole-brain bauplan based on synaptic annotation remains a demanding effort. As an alternative, we present here an approach to extract an approximated brain connectome starting from libraries of single cell anatomical reconstructions aligned on the same anatomical reference brain. Our approach relies on the identification of neurite terminal nodes based on Strahler numbering of the cell morphology, and the adoption of a proximity range criterion, so as to infer, in the absence of synapse information, approximated brain network architectures. As an initial benchmark we used information theory metrics to confront our approach against a synaptically-annotated EM dataset of *Drosophila melanogaster* hemibrain. The comparison with our approach revealed a general agreement in the organization of the extracted connectivity modules measured in terms of Normalized Mutual Information, Adjusted Rand Index and Pearson correlation. Moreover, we show that the modules identified, along with their organization, can capture known circuit motives. We then applied this approach to a light microscopy dataset of the zebrafish larval brain composed of about 3000 neuronal skeletonizations. We show that the approximated connectome and the resulting modular organization is capable of capturing specific and topographically organized connection patterns as well as known functional circuit architectures. In conclusion, we present a scalable, from-circuit-to-brain range, approach to reveal approximated neuronal architectures supporting brain mechanisms, potentially suitable for hypothesis generation and for guiding the exploration of and integration with EM connectomes.

**Author Summary:** Understanding how the brain works requires detailed maps of its neural connections, known as connectomes. While advanced techniques like electron microscopy (EM) can map these connections at the level of individual synapses, they are time consuming and resource intensive, limiting their use. As an alternative, we developed a computational method that approximates brain connectivity using existing datasets of neuron shapes (morphologies) without requiring synaptic annotations. Our approach identifies potential connections between neurons based on the proximity of their terminal branches regions where synapses are likely to form within a shared 3D reference brain. We validated our method using a high-resolution EM connectome of the fruit fly brain, demonstrating that it captures broad organizational patterns, such as clusters of densely interconnected neurons, despite moderate agreement at the synaptic level. Applying the method to a light microscopy dataset of the zebrafish larva brain (∼3,000 neurons), we successfully reconstructed large-scale networks that recapitulated known functional circuits. This approach offers a scalable way to extract brain-wide connectivity principles from existing datasets, bridging the gap between cellular anatomy and circuit function. It can guide targeted experiments and complement future EM studies, making neuroscience data more accessible for hypothesis generation.

## Introduction

The understanding of the brain mechanisms takes advantage of the continuous refinement of molecular, hardware, and software approaches. These methods allow reconstructing, in suitable model organisms, the brain dynamics from synapse level to brain-wide scales ^1–5^. Current optical and electrophysiological methods represent invaluable tools for studying the neuronal dynamics associated with sensory information, the fine-tuning of motor programs, and spatiotemporal patterns characterizing higher-level brain activity ^6–8^. The resulting dataset becomes fundamental building blocks for the formulation, and the possible falsification, of circuit models describing the functional working mechanisms ^9–11^. However, even under optimal conditions, one faces a typical scenario where different models, despite their intrinsic architectural differences, number of parameters, and complexity, perform similarly well in capturing the same circuit dynamics ^12,13^. With the aim of better constraining this kind of problem, one possible strategy is offered by the availability of the anatomical information characterizing the circuit wiring diagram supporting a certain functional mechanism. Then, it is convenient to try to refine a circuit model considering the wiring properties of the underlying circuit. For this purpose, one can take advantage of dense anatomical reconstructions of large brain parcels based on Electron Microscopy (EM) ^14–18^, which provides sufficient resolution levels to characterize the synaptic contacts and possibly the neurotransmitters involved between neuronal pre- and post-synaptic partners, while covering a large field of view. Retrieving a whole-brain connectome, even in accessible organisms, demands extensive acquisition time. This includes supervised or semi-supervised tracing, cell segmentation, synapse annotation, and thorough manual proofreading. Today, even with the support of AI-based algorithms, this is still a time-demanding process that limits its applicability to a few samples and requires computational strategies to minimize human workload ^15,19^. Once available the information on the neuronal connectivity schemes i.e. the organization of the input-output contacts characterizing the cells involved, one can obtain an adjacency matrix representing a connectome and identify the circuit wiring diagrams. Because of the amount and the complexity of the data one has to face, this type of anatomical reconstruction frequently focuses on targeted circuit motifs, specific neuronal subpopulations, or particular brain regions. Currently, even if there are several reported whole-brain EM datasets ^15,16,18,20–22^, partial or whole-brain connectomes that provide skeletonization of neurons along with detailed annotations of their synaptic contacts are rather limited. This applies also for circuit wiring reconstruction approaches relying on techniques other than EM. Using light microscopy, with traditional configurations or leveraging super-resolution/expansion microscopy techniques, in combination with fluorescence labelling for reconstructing synaptic partners, it has been shown to reliably capture the connectivity scheme in different circuits or brain regions ^23–26^. Targeting two different fluorophores or leveraging single fluorophore reconstitution to pre- and post-synaptic sites respectively is another of the available methods to map effective connectivity ^27–30^. Using a different approach researchers have used confocal microscopy in combination with genetic labelling to visualize and manually reconstruct ∼16000 neurons of D. melanogaster and to identify based on hierarchical clustering a number of fundamental local units ^31,32^. Unraveling the anatomical organization of brain-wide networks requires suitable datasets, with a sufficient sampling of the cellular morphologies, coverage across different regions of the brain and ideally based on the evaluation of a large number of samples. To complement the methods currently available, here, we present a computational approach to generate an approximated connectome starting from an anatomical dataset with neuronal cell skeletonizations, belonging to or co-registered in the same reference space, where the actual synaptic information is not accessible or not yet available ^33,34^. Similarly to other proposed approaches ^33–35^, the purpose is to use already existing neuromorphological datasets (either EM or LM) of brain organisms such as *Drosophila* and zebrafish, and present a computational approach to extract approximated basic wiring principles of the brain circuits. For example, the approach proposed here can be applied on mesoscale atlases based on optical microscopy acquisitions, as well as electron microscopy datasets not yet fully annotated at the detail level corresponding to the neuronal input-output features ^36–41^. For this type of dataset, here, we devised an approach to infer brain network organization by assigning cell-to-cell connections, which are subsequently translated into adjacency matrices, representing the brain wide cell resolution connectivity scheme. Following its evaluation on EM-based ground truth, we applied this approach to a zebrafish single-cell dataset, which comprises more than three thousand neuronal reconstructions.

## Results

### 1. Constructing approximated connectomes based on Strahler numbering and spatial proximity

For the cases where actual synaptic information is unavailable, we devised an approach to infer an approximated brain network structure by assigning cell-to-cell connections on the basis of a proximity rule between the nodes along the reconstructed cell skeleton. We worked along a research line similar to previous studies^34^ (see the Methods section). In our case, we began by analyzing the cell anatomy and calculated the Strahler’s number (SN) for each point along the cell skeleton. The SN is a parameter that indicates the relative hierarchical level of a point within the branching architecture of the cell ^41,42^, with the soma corresponding to the highest SN and terminal points along the neurites with the lowest (Figure 1A). Previous studies have shown that a greater proportion of synaptic partners is associated with terminal points (SN = 1) ^43–46^. We considered terminal nodes (i.e. neurites segments with SN = 1) as potential sites of connection between different cells, if they fell within a defined proximity range with respect to nodes belonging to other cells ^34^. For a typical light microscopy dataset, considering the intrinsic lateral resolution of the acquisition process (∼1 *μ*m), the typical errors in the anatomical reconstruction (∼1 *μ*m), and intrinsic variability in the anatomical co-registration procedures (∼5 *μ*m with respect to the 4.38 *μ*m reported in the original manuscript), we adopted a proximity ranging between 0 and 5 *μ*m (Figure 1B).

**Figure 1.**
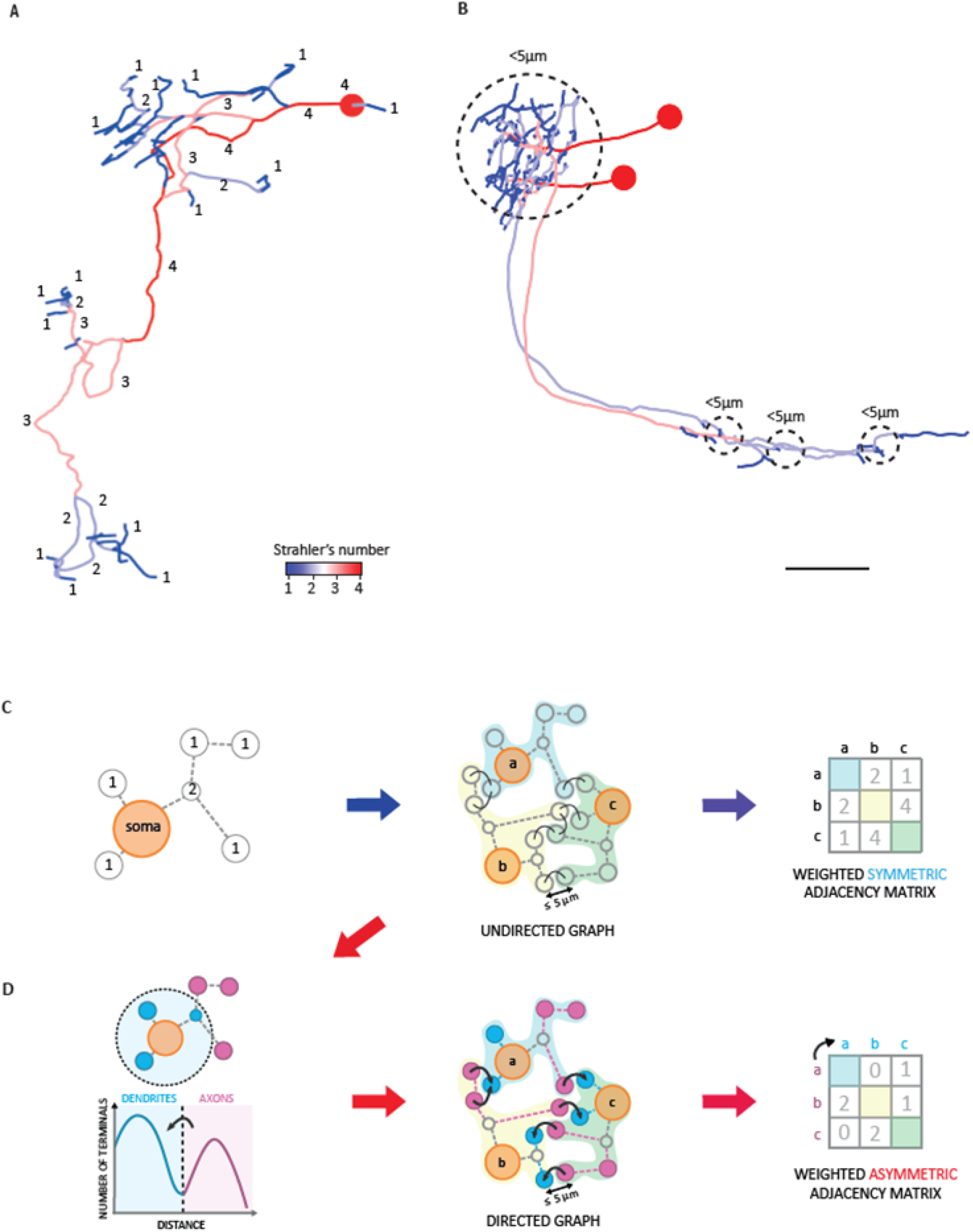
**A)** Strahler Number (SN) annotation of the neuronal anatomy. Numbers indicate the relative level of exemplar nodes in the bifurcation hierarchy of the cell neurites and the color map represents the corresponding SN value. **B)** Identification of the putative synaptic contacts. Based on the proximity range adopted (0-5 *μ*m in this case), neurite terminal nodes, i.e. with a SN equal to 1, are identified. **C)** Construction of the undirected graph. Based on the proximity range, input-output pairs are annotated and assigned to every neuron (a,b,c,…), so as to distill a weighted symmetric adjacency matrix underlying an undirected graph. **D)** Construction of the directed graph. A directed graph is built based on the subcellular compartmentalization of the input vs the output nodes with respect to the distance from the cell soma.

As reconstructing the connectome relies on the determination of the adjacency matrix *A*_*i,j*_, we envisaged two different possible scenarios. In the first one, we assumed no subcellular segregations for the input and output nodes of a neuron, resulting in a symmetric adjacency matrix 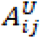 corresponding to an undirected connectome (Figure 1C). In the second one, as most of the neuronal reconstructions available are characterized by a bimodal spatial distribution of the neuronal nodes^31^ (Supplementary Figure 1), we assign for each cell a separation in two putative components: input dendritic points in the regions proximal to the soma, and output axonal points in more distal areas. This allowed us to constrain the connectome according to an axon-to-dendrite synaptic scheme. In this latter case, we obtain an asymmetric adjacency matrix 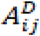 representing a directed connectome (Figure 1D).

### 2. Evaluating the connectome reconstruction strategy against an EM-based ground truth

To assess the actual applicability of the approach and its capability to capture brain circuit organization patterns, we took advantage of a Drosophila melanogaster hemibrain EM dataset characterized by a large fraction of the cells (approximately 25,000 cells, corresponding to about 22% of the estimated total) with input-output synaptic annotation ^47^. We initially explored the structural organization of brain-wide networks, using the Leiden algorithm ^48^, a method based on modularity maximization to identify clusters of densely connected neurons within the full dataset (see Method section for details). Applying this method on the synapse-annotated EM dataset, we uncovered a hierarchical organization composed of eight main modules (module 1-8), each accounting on average for about three thousand cells. Re-iteration of the method at the single module level led to the identification of a number of submodules ranging from four to nine (Figure 2A, Supplementary Figures 10-17), associated with each main module. For each main module identified on the EM-annotated dataset, we then calculated again the adjacency matrix and the modularity organization, but this time without considering the available synapse annotation, but relying solely on the Strahler numbering of the cell skeleton and on the proximity range definition to build an directed graph (see Methods section). The pairs of adjacency matrices obtained, one based on the synapse annotation (EM-based) and one relying on the Strahler numbering (Strahler-based), present Pearson correlation coefficients ranging from 0.05 to 0.215 always with p-value smaller than 0.001 (see Methods), with minimal variations considering the 1 *μ*m and 5 *μ*m proximity range (0.146 ± 0.053 and 0.14 ± 0.052, average ± SD, respectively). In order to evaluate the capability of the approximated approach to actually capture the expected wiring, we calculated for each of the main modules identified in the EM-based analysis the Adjusted Rand Index (ARI) ^49–51^ based on the cell grouping associated with the organization of the corresponding submodules. This metric is frequently used for comparing different partitions or clustering patterns and returns a value between -1 and 1 to quantify the level of overlap in the contingency matrix. For the 1 *μ*m proximity range, we obtained an average ARI of 0.36 ± 0.17 (average ± SD, covering a range between 0.04 and 0.6), pointing to a general overlap of the Strahler-based approach with the ground truth sub-communities. A similar trend resulted for the 5 *μ*m proximity range, 0.38 ± 0.17 (average ± SD, Figure 2B, covering a range between 0.04 and 0.6). To compare the community detection accuracy between the two datasets, we also computed the Normalized Mutual Information (NMI) ^52,53^. This metric returns a parameter between 0 and 1, depending on the level of overlap between the communities using adjacency matrices and modularity analysis obtained with the two approaches. NMI ranged between 0.1 and 0.55 (0.38 ± 0.17 average ± SD) and between 0.1 and 0.56 (0.38 ± 0.17 average ± SD) for 1 *μ*m and 5 *μ*m, respectively (Figure 2B). Then, to corroborate this outcome, we evaluated the significance of the calculated ARI and NMI scores, with a comparison against two types of random null models. We calculated the Z-score for NMI and ARI between the Strahler-based connectome modules and the EM-based connectome modules, focusing here on two modules in particular, module 6 and module 8, those characterized by the lowest and the highest, ARI and NMI values. We constructed 100 random networks for the two Strahler-based connectome modules either by randomizing the links or the weights of the links between the neurons (see Methods). Then, we quantified the p-value by counting the number of random networks that had their own z-scores (for ARI and NMI) higher than that one of the corresponding Strahler-based modules ^54^. For each module we found *p* < 0.05 highlighting that the Strahler-based model can capture connectivity patterns in the networks better than models with the same number of nodes, but different links or link strength organizations. Similar analysis has also been done for Pearson correlation, obtaining the same conclusion as ARI and NMI, as shown in Supplementary Figure 18. To further validate our results, we compared the Pearson correlation between the inferred and true connectivity against 100 degree-preserving random null models. For Module 6, the Pearson correlation between true and inferred connectivity was 0.214, significantly higher than the mean correlation of 0.0524 obtained from randomized networks, yielding a z-score of 31.94 (with randomized means near zero and a maximum z-score of 4.14). Similarly, for Module 8, the Pearson correlation was 0.144 compared to a randomized mean of 0.0165, with a z-score of 31.82 (randomized means near zero, maximum z-score 5.97). These results demonstrate that while the absolute Pearson correlations may appear modest, they are statistically robust when compared to randomized networks, which perform substantially worse. This confirms that our inferred connectivity captures meaningful structure beyond chance.

**Figure 2.**
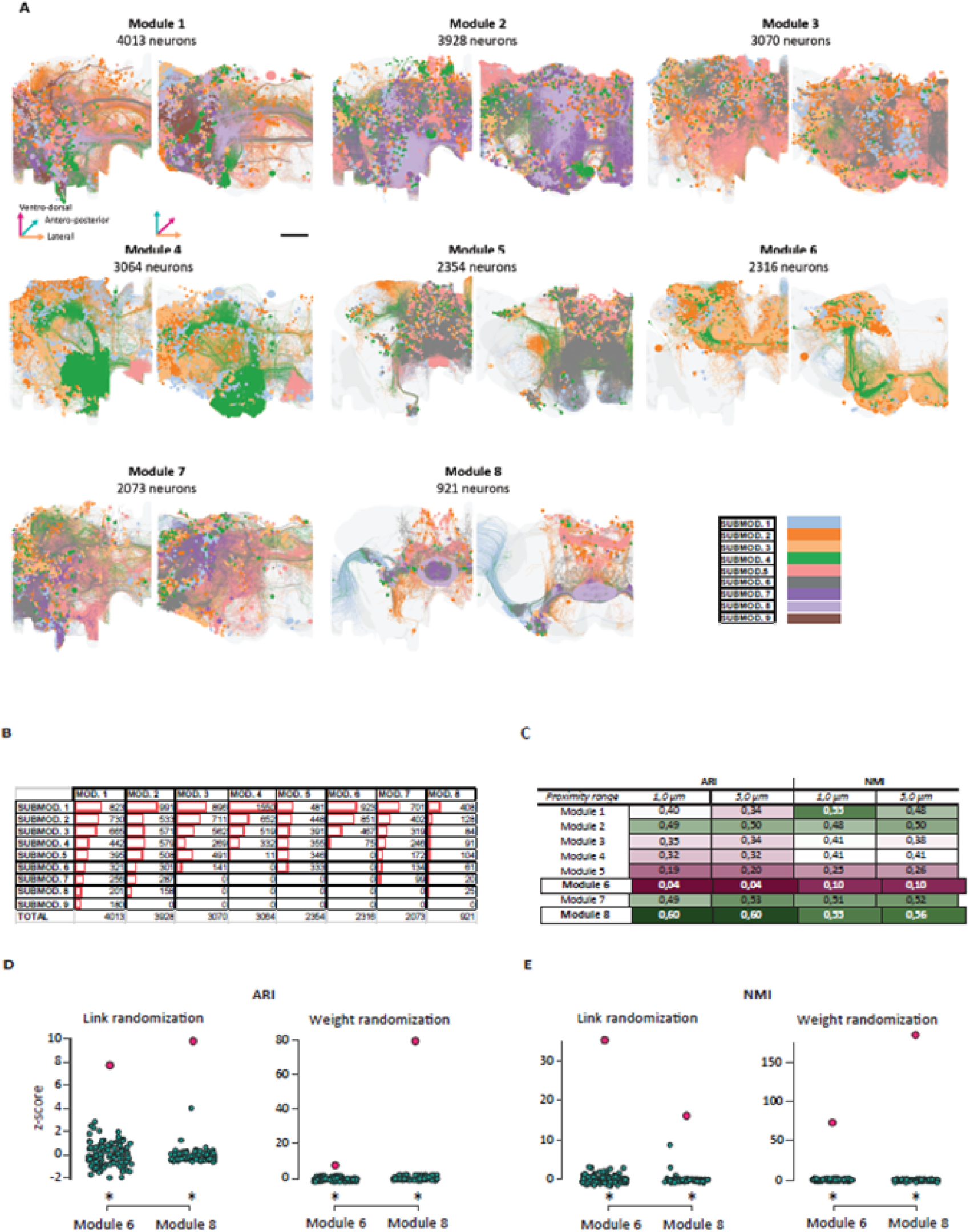
Comparison with EM-based ground truth and null models. **A)** Decomposition of the EM Drosophila dataset in higher order modules and corresponding submodules obtained with the second iteration of the algorithm. Colors are assigned according to the submodules.. **B)** Corresponding submodule organizations associated with the identified higher order modules. **C)** ARI and NMI computed between the EM (“ground truth”) and Strahler reconstructed network at different proximity ranges. **D**,**E)** Z-scores computed over the ARI (panel **C**) and NMI (panel **D**) between Module 6 (or 8) of EM and Strahler reconstructed network (red point) and z-scores computed over the same mentioned similarity measures between EM and 100 random null models of Strahler reconstructed network (green points). The larger the gaps between the red point and the green points, the farther is the Strahler based networks from random behavior and closer is to the ground truth (EM based networks). *: p<0.05.

While the connectivity modules identified with our approach show significant similarity to EM-based ground truth (p<0.05), on the other side the correlations (Pearson: 0.05–0.215), Adjusted Rand Index (ARI: 0.04–0.6), and Normalized Mutual Information (NMI: 0.1–0.56) indicate moderate agreement. This underscores the method’s utility for approximating large-scale connectivity patterns, though with inherent limitations in resolving fine synaptic specificity.

#### 2.1 Degree Distribution Comparison

To explore the relationship between proximity range selection and degree distribution similarity, we conducted an analysis comparing the degree distributions of the EM-based Drosophila Hemibrain connectome with those reconstructed using the Strahler-based method across proximity ranges of 1 *μ*m and 5 *μ*m for all modules (1–8). Visual inspection of the distributions suggested that a proximity range of 1 *μ*m or lower may yield better overlap with the ground-truth EM distributions. To further investigate the utility of degree distribution overlap as a criterion for parameter selection, we focused on Module 8 (921 neurons) due to its tractable size. We reconstructed the module using proximity ranges of [0.2, 0.4, 0.6, 0.8, 1.0, 1.2] *μ*m and compared the resulting degree distributions to the EM ground truth (Supplementary Figure 9). Overlap was quantified using the Kolmogorov-Smirnov (KS) test and Jensen-Shannon (JS) divergence (a symmetric alternative to KL divergence), implemented via Python’s *scipy*.*stats* and *scipy*.*spatial* libraries, respectively. The key observations from performing the KS Test found that for out-degree distributions, the KS statistic reached a minimum (∼0.27, *p* < 0.01) at 0.4 *μ*m, indicating closer alignment with the EM distribution at smaller proximity ranges. In contrast, in-degree distributions showed slower variation with proximity range, suggesting less sensitivity to this parameter. Similarly, JS Divergence comparison showed that the JS measure for out-degree distributions was minimized (∼0.444, *p* < 0.01) at ∼0.6 *μ*m, further supporting the trend observed with the KS test. In-degree distributions again exhibited weaker dependence on proximity range.

#### 2.2 Performance evaluation of strahler-based network reconstruction using False positive and negative analysis

We evaluated the performance of predicted connectivity graphs which we refer to in our work as Strahler-based method against ground-truth graphs derived from EM data across eight modules in the *Drosophila* brain. These performance metrics are defined as follows: For every edge in the union of predicted and true graphs, we classified it as a *true positive* (TP, if present in both), *false positive* (FP, predicted but absent in ground truth), or *false negative* (FN, absent in prediction but present in ground truth). *True negatives* (TN) were computed as all possible edges (given the node set) minus TP, FP, and FN. From these counts, we calculated standard metrics: False positive rate (FPR=FP/[FP+TN], False negative rate (FNR=FN/[FN+TP]) and accuracy (proportion of correctly predicted edges).

First we evaluated the performance between strahler-based predicted graph and ground truth across all modules by keeping proximity range 1*μ*m fixed and the results are summarized in Supplementary Table 1. We found that Module 2 achieved the highest accuracy (0.9771) and lowest false positive rate (FPR: 0.0058), suggesting robust prediction for this subcircuit. Module 5 had the lowest accuracy (0.9306) and highest false negative rate (FNR: 0.9813), indicating under prediction of true edges. We also found that Module 8 showed intermediate accuracy (0.9084) but a balanced FPR/FNR (0.0063/0.9365), reflecting a trade-off between sensitivity and specificity.

We then delved further into the performance evaluation of Module 8 by observing the impact of proximity range at finer proximity intervals (0.2–1.2*μ*m), the results of which have been summarized in Supplementary Table 2. We observed a clear trend i.e. accuracy values declined slightly from 0.9103 (0.2*μ*m) to 0.9079 (1.2*μ*m), reflecting a trade-off between edge inclusion and correctness. On the side of error rates, we found that FPR increased monotonically (0.0028 to 0.0070) as the proximity range increased, capturing more potential false edges however for FNR where it improved (0.9503 to 0.9347), indicating better recovery of true edges at larger proximities.

However, overall we find certain observations and acknowledge certain limitations from this performance evaluation. There is a trade-off between accuracy and FNR where higher accuracy modules (for e.g., Module 2) still exhibited substantial FNRs (0.9130), highlighting systematic underprediction. There can also be impact of sparsity of connections in the predicted model or even “over connection” in ground truth data leading to low FPRs (e.g., 0.0058 for Module 2) which may suggest conservative edge prediction, while high FNRs reveal missed biological connections. The stringent requirements of binarizing the edges of true and predicted graphs before computing the metrics, without the consideration of weights and thresholding them can also affect their performance evaluation. Finally, we find that performance metrics differ significantly across modules, implying that prediction reliability depends on subcircuit properties (e.g., size, density). And the two parameters such as proximity range tuning (as seen from Module 8, Supplementary Table 2) along with proper thresholding to binarize edge weights of true and predicted graphs before computing metrics can further improve the overall structural similarity between the ground truth and strahler predicted graphs.

#### 2.3 Modularity highlights

Having established that the approximated connectome effectively captures connectivity patterns, we proceeded to assess whether the modules identified with our approach hold functional significance. Our analysis focused on a single higher-order module (module 8) which has the highest ARI and NMI values. This module is primarily associated with the central complex (CX), and more specifically with the ellipsoid body (EB) network (Supplementary Figure 17). Decomposing this module, with a second re-iteration of the Leiden algorithm, revealed a set of lower-order components that can be grouped into three distinct networks, each characterized by unique spatial segregation patterns possibly reflecting specialized functional roles. The first network (Supplementary Figure 17, submodule 8.1) comprises neurons that project in the anterior optic tubercle (AOTU) and the early visual pathway. These are primarily medulla columnar neurons that receive visual inputs and convey them to the bulb (BU)^55^. The second network (Figure 3B, submodules 8.3, 8.4, 8.6, 8.7, 8.8) consists mainly of tangential neurons that transmit signals from the BU to the ellipsoid body (EB) forming a series of concentric lamination domains^55,56^. The third network (Supplementary Figure 17, submodules 8.2, 8.5) comprises columnar neurons projecting to the protocerebrum (PB) and the EB involved in the interpretation of spatial information^56,57^. Interestingly the approach identified a spatial organization in this columnar projection stream supporting the space representation^58^.

**Figure 3.**
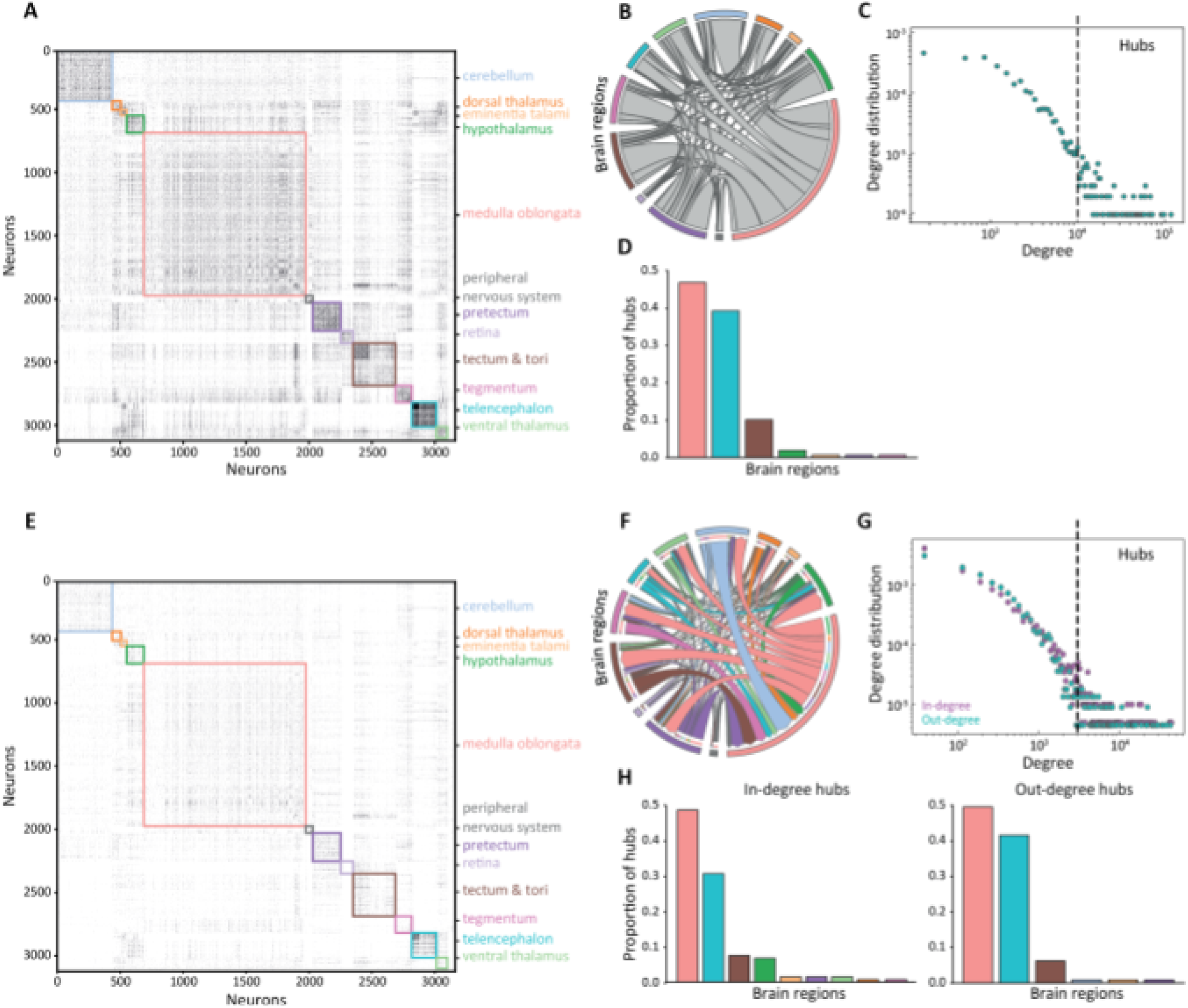
Topological properties of undirected and directed networks. **A, E)** Binarized adjacency matrixes showing connections in the undirected (A) and directed (E) networks with neurons sorted by brain regions in which the soma is located. The matrix is binarized for visualization purposes. **B, F)** Chord plots representing inter-regions connections for the undirected (B) network and incoming – outgoing inter-regions connection for the directed network (F). Labels for the colors are reported in A and E. **C, G)** Weighted degree distributions for the undirected (C) and directed (G) network. Doshed lines depict the threshold above which neurons are defined as hubs. **D, H)** Bar plots representing the distribution of hubs in the brain regions for the undirected (D) and directed (H) networks. Labels for the colors ore reported in A and E.

### 3. Constructing an approximated connectome of the zebrafish larval brain

Aiming to build an approximated connectome for the zebrafish larval brain, we took advantage of an available dataset with anatomical reconstructions of individual neurons obtained by means of optical microscopy z-stack acquisitions co-registered on the same reference brain ^40^ (Figure 1A). Applying the proposed method, we retrieved two connectome scenarios showing a rather articulated scheme of neuronal interactions between the different regions of the brain (Figure 3B, Figure 3F). From the obtained adjacency matrices,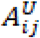, and 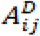 (Figure 3A and Figure 3B), we extracted a set of graph parameters. In general, being the non-zero entries of the directed matrix a subset of the undirected ones, 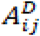 resulted sparser than 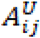 . The analysis of the undirected graph,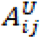, revealed the existence of 23 connected subgraphs, with a Giant Component (GC) accounting for 99% of the edges of the original graph and a set of small, poorly populated disconnected subgraphs (accounting for 32 out of total 3163 cells), that have been neglected from further analyses. The GC presents a network density of 0.0397, pointing to a rather limited connectivity, a small weighted clustering coefficient of 0.002 and an average shortest path length of 3.792. The undirected graph shows a heterogeneous degree distribution, with most of the nodes characterized by a relatively small number of connections (<10^3^) and fewer ones – hubs-like - with high degrees (>10^4^) (Figure 3C). Interestingly, for low-degree nodes, the distribution shows a flat trend, suggesting deviations from the power-law behavior. An assortativity analysis gives a correlation coefficient *β* =0.345, meaning that the graph is assortative and that nodes tend to connect with those of a similar degree i.e., small-degree nodes to small-degree nodes and hubs to hubs. We mapped hubs to the corresponding anatomical regions normalizing for the relative number of cells; it appears that the majority belongs to Medulla Oblungata (Figure 3D - Pink), followed by the Telencephalon (Figure3D - Cyan) and the Tectum Opticum (Figure 3D - Brown). On the other hand, regarding the weighted directed graph, 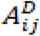 (Figure 3E), we found a low clustering coefficient (0.0005) and a path length of 6.10. While the in- and out-degree distributions resemble the general heterogeneity of the undirected scenario, the low-degree region did not show a corresponding saturation (Figure 3G).

### 4. Analysis of brain-wide modules from the approximated zebrafish larvae connectome

One of the most appealing uses of a brain wide connectome is unravelling possible network architecture and circuit organization motives that characterize an organism brain. We then considered using the Leiden algorithm, a network community discovery approach based on modularity optimization, i.e. the Leiden algorithm^48^, to analyze the network organization of the zebrafish connectome. Running the algorithm 10,000 times on the undirected GC typically resulted in the stable and robust identification of 12 neuronal modules, with small deviations in the assignment of a few nodes from trial to trial (<0.5%). However, only eight of the identified modules are prominent and well populated, the remaining containing an extremely limited number of nodes (4 out of 3116 and 14 out of 3131 in total for directed and undirected GC graphs) (Figure 4A). To characterize the resulting networks, we first mapped all identified undirected modules (U) on the brain regions containing the body of neurons belonging to each corresponding module (Figure 4B). In general, all modules present rather distributed spatial patterns, covering a number of brain regions in a non-exclusive manner. In addition, most of the brain regions in this analysis, although to different extents, participate in more than one module (Figure 4C). Similar to the undirected connectome, the analysis of the directed connectome revealed the presence of seven main modules (D) (Figure 4E-G), showing a substantial overlap, beside two directed modules. D2 includes two undirected modules, U4 and U6, and D1 combines U1 with a fraction of U3 (Figure 4D, H). This substantial overlap between the two modularity analyses, directed and undirected cases, points to the possibility that the two corresponding connectomes generated in this way rely to a large extent on a common backbone represented by a directed graph and axon-dendrite communication as reported for other datasets with synapse annotation^21^.

**Figure 4.**
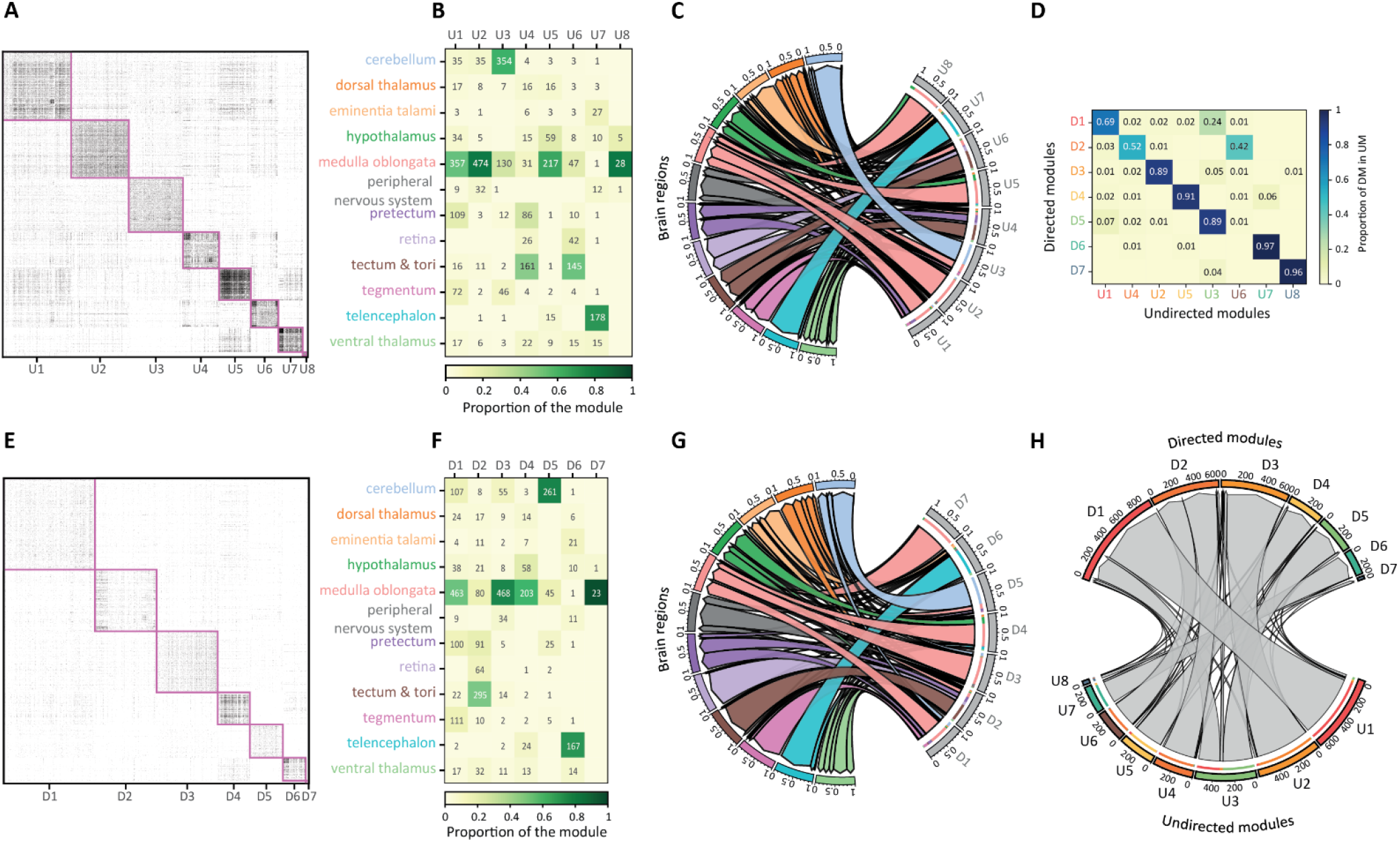
Distribution of the modules in the different regions for the undirected and directed networks. **A,E)** Adjacency matrices recapitulating neuronal connections in the undirected (A) and directed (E) networks sorted by the identified modules. The matrix is binarized for visualization purposes only. **B,F)** Distribution of the modules across the brain regions for the undirected (B) and directed (F) network. Number refers to neurons belonging to a certain module and with the soma located in the regions. Matrix cells are color-coded based on their proportion within the module. U: undirected module, D: directed module. **C,G)** Proportion of the distribution of the modules across the brain regions for the undirected (C) and the directed (G) networks. Labels for the colors are reported in B and F. **D)** Proportion of neurons of the undirected network modules present in the directed network modules. **H)** Number of neurons of the undirected network modules present in the directed network modules. Labels for the colors are reported in D.

### 5. Hierarchical anatomical organization of the modules reflects circuit functional role

To better capture the 3D anatomical organization of the 8 identified modules in the undirected graph, we represented the wiring diagrams corresponding to each module in the anatomical reference space (Figure 5). In some of the cases, the identified modules correspond to previously described brain-wide circuits underlying information processing or motor control functions, like the visual network module ^41^ (Figure 5D and Figure 5F) and the cerebellar network module (Figure 5C). To unravel finer structures in the identified modules, we re-iterated the Leiden algorithm in each main module. This approach, where each module is considered an independent subnetwork, led to the finding of a variable number of subcomponents, typically ranging between five and eight (Figure 5 and Supplementary Figures 2 - 7). These submodule patterns, even with the limitations deriving from the neuronal contact approximation employed here, captured an additional level of the circuit organization, involving features of spatial organization of the neuronal populations across the brain.

**Figure 5.**
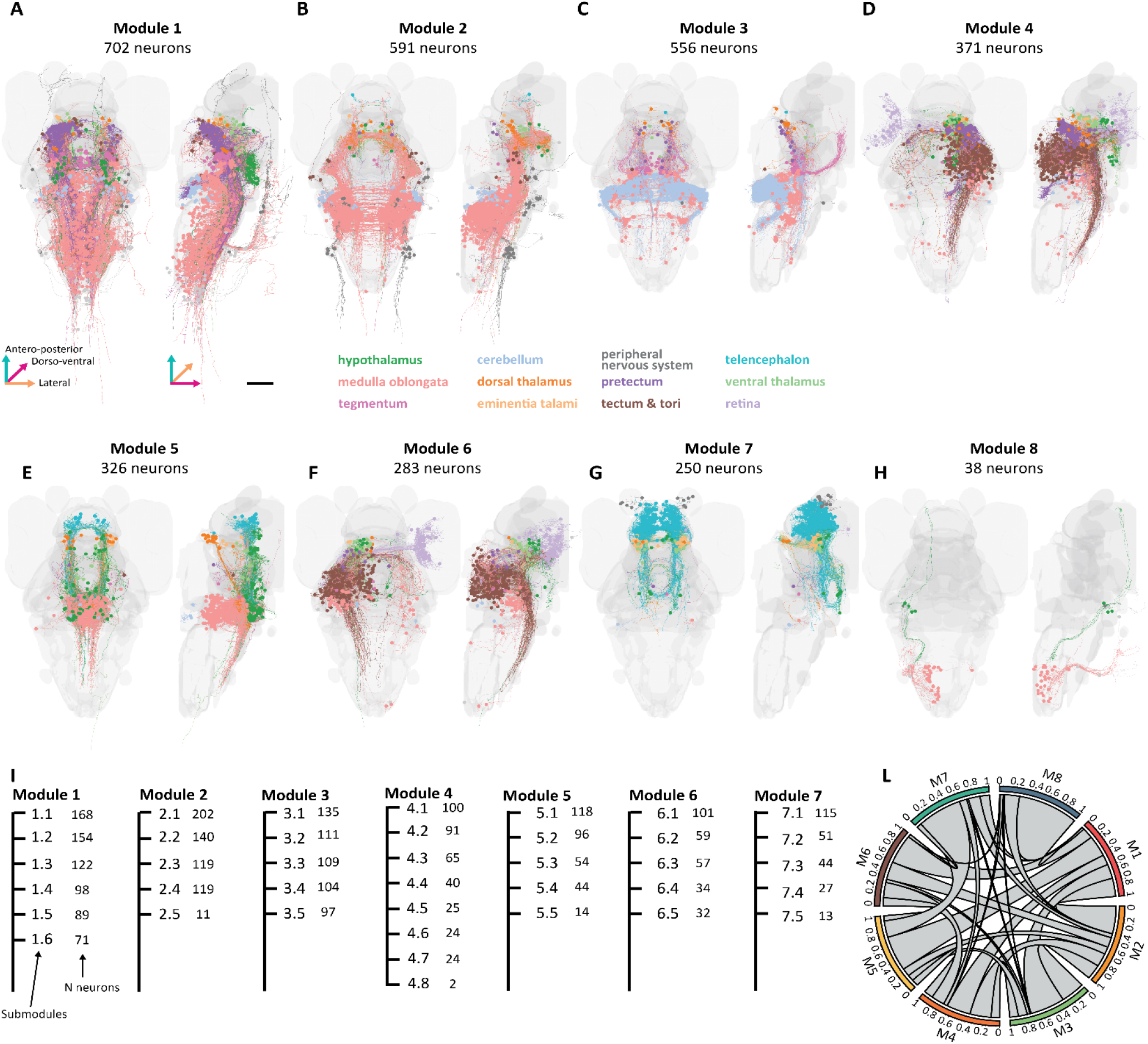
Modules of the zebra fish brain. **A-H)** Anatomical representation of the eight high-order modules with the corresponding submodules found in the zebrafish dataset overlaid on the brain silhouette. Colors are assigned to the cells according to the soma position in the different brain regions. Scale bar =100 *μ*m. **I)** Tree diagram reporting the structure of the zebrafish network with the organization of the submodules for the different modules. **L)** Distribution of connections between the different modules.

First, we considered the modules including the organization of the Retinal Ganglion Cells (RGCs), projections to the tectum (modules 4 and 6). The analysis of the relationship of the RGC body position in the retina and the connected PeriVentricular Neurons (PVNs) in the contralateral tectum showed a retinotopic organization in submodules 4.2 and 4.6 (right tectum, Figure 5A), and in the sub modules 6.1 and 6.2 (left tectum) (Figure 5B). Indeed, these submodules show projections from RGC cells with soma located in the caudal part of the retina to connect with PVNs in the rostral part of the contralateral periventricular region at the level of the tectum and, conversely, for the RGC connections involving the rostral part of the retina and the caudal part of the contralateral retino-recipient area. Similarly, our analysis revealed an articulated spatial pattern of connections involving different brain areas associated with the module 3, representing a pretectal-cerebellar network (Figure 6C). The connections formed between neurons located in the medulla oblongata, cerebellum, and tegmentum (Figure 6C1, submodules 3.3 and 3.4) resemble those associated with vestibulo-ocular reflex (VOR) ^59^. In this case, the ascending and ascending/descending tangential neurons are located in the medial octavolateral nucleus (MON) of the medulla (Figure 6C1, submodules 3.3 and 3.4), the recipient of utricular inputs. These neurons project to the oculomotor nucleus (nIII) where extraocular motor neurons mediate gaze stabilization ^59^. Furthermore, connections between neurons located in the cerebellum have been shown to be involved in VOR ^60,61^. The cerebellum displays a spatially segregated pattern of distinct cells, with lateral Purkinje cells projecting to the MON (Figure 6C2, submodule 3.2) and eurydendroid cells located in the middle with less specific projections ^40,62^ (Figure 6C2, submodules 3.1 and 3.5). Even though our approach highlighted the existence of principal networks supported by modules of neurons characterized by a relatively higher connection degree, it is clear that the different identified subnetworks show mutual intermodule connection schemes. In order to quantify this aspect, we evaluated the links formed across different modules (see Supplementary Figure 8). This analysis showed a rather heterogeneous scenario, with some modules characterized by a relevant number of intermodular connections, while others with a very limited number of intermodules links.

**Figure 6.**
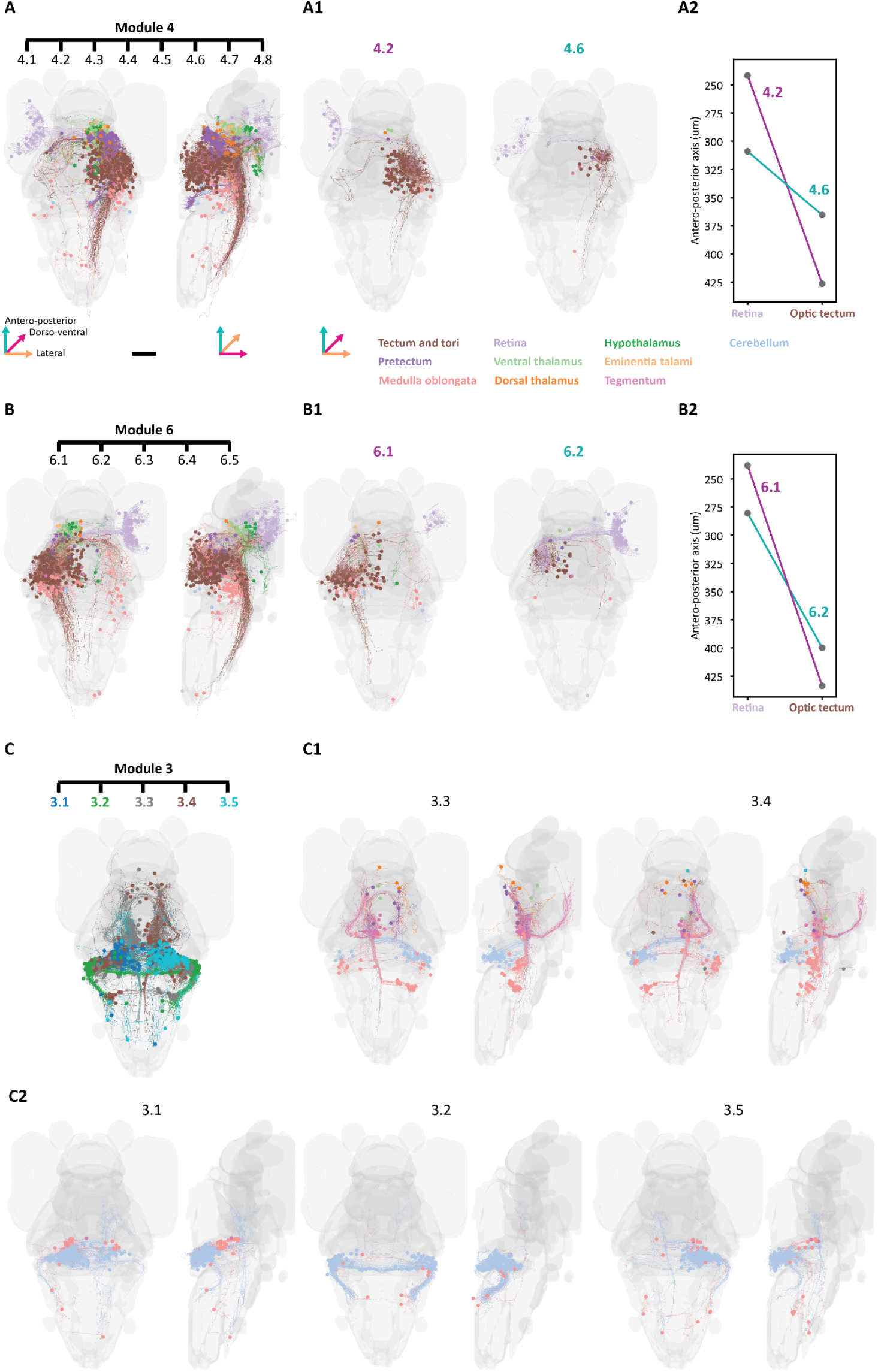
Functional description of the modules. **A)** Anatomical representation of the high-order module U4. Colors are assigned according to the soma distribution in the brain regions. Scale bar = 100 *μ*m. **A1)** Sub-modules associated with the right Tectum opticum involving PeriVentricular Neurons and Retinal Ganglion Cells, highlighting two different sub-populations of neurons (4.2 caudal tectum, medial tectum). **A2)** Median of the distributions along the antero-posterior axis of retina and optic tectum neurons of modules 4.2 (magenta) and 4.6 (teal). **B)** Anatomical representation of the high-order module U6. Colors are assigned according to the soma distribution in the brain regions. **B1)** Sub-modules associated with the left Tectum opticum involving PeriVentricular Neurons and Retinal Ganglion Cells, highlighting two different sub-populations of neurons (6.1 caudal tectum, 6.2 medial tectum). **B2)** Median of the distributions along the antero-posterior axis of retina and optic tectum neurons of modules 6.1 (magenta) and 6.2 (teal). **C)** Anatomical representation of the high-order module U3 as composed by the five lower-order modules (3.1-3.5). Colors are assigned according to the lower-order modules. MON: medial octavolateral nucleus, nIII: oculomotor nucleus. **C1-C2)** Anatomical representation of the individual lower-order modules. Colors are assigned according to the location of the soma. Scale bar = 100 *μ*m.

## DISCUSSION

The development of methods to infer brain-wide connectivity in the absence of synaptic-resolution data could represent an important step toward leveraging existing neuroanatomical datasets for network neuroscience. Here, we present a computational framework that combines Strahler order analysis and spatial proximity criteria to approximate connectomes from libraries of co-registered neuronal reconstructions. By validating our approach against a *Drosophila* EM dataset and applying it to a zebrafish larval brain atlas, we demonstrate its ability to recover first approximate organizational principles of brain networks, albeit with inherent limitations in resolving synaptic specificity.

The proposed approach infers an approximated cell-to-cell contact structure based on a spatial proximity rule and can be applied to any dataset with single cell reconstructions where synapse direct annotations are unavailable or incomplete. This typically is the case for anatomical datasets derived from sparsely labeled light microscopy, such as those acquired through laser scanning or light-sheet microscopy^36–40^. In such scenarios, even with the addition of synaptic tagging strategies involving pre- and post-synaptic partner labeling^63,64^, the information retrieved is often partial or biased—potentially omitting important circuit elements. Conversely, high-resolution reconstructions from dense tissue datasets, such as those obtained via serial block-face electron microscopy, offer synaptic-level detail but produce volumes of data that remain difficult to process at scale^47^. Although automated reconstruction of cell morphology and their manual validation could become possible for small datasets, a complete synapse annotation may still represent a prohibitive challenge. These constraints underscore the need for complementary strategies that use available morphological skeletons to infer approximate connectivity across entire brains.

Our framework addresses this need by estimating potential connections between neurons based on proximity thresholds applied to their reconstructed morphologies. While this cannot achieve synaptic specificity, it captures essential aspects of network organization, offering a mesoscale perspective on brain architecture. The approach is not meant to replace high-resolution connectomics in the characterization of the microcircuit structure^65^ but rather to complement it by seeking to understand the general organization of brain networks ^66^.

We validated our approach using the synapse-annotated connectome of the *Drosophila melanogaster* hemibrain, containing 21,739 connected neurons^47^. Through modularity analysis, we uncovered a hierarchical organization comprising eight main modules, each with several submodules. When comparing the adjacency matrices generated from synapse annotations (EM-based) and those derived from the spatial proximity rule, we observed significant correlations with respect to two null models where links or weights have been re-shuffled. Thus, confirming that the spatial proximity rule can successfully capture broad organization principles of brain connectivity.

Having established its validity, we next applied the method to a mesoscale zebrafish (*Danio rerio*) brain atlas comprising approximately 3,200 single-neuron reconstructions lacking synaptic annotations. This analysis revealed a set of neuronal modules—discrete communities with relatively high internal connectivity—that effectively outline a scaffold or “backbone” of the zebrafish brain’s wiring diagram. With this approach, we have obtained a decomposition of the connectomes of two phylogenetically distant organisms on a set of basic components that form a sort of neuronal backbone of the brain wiring diagram. For some of the identified modules, it is possible to associate a functional role, either based on known anatomical characterizations of the circuits involved, or on the evidence provided by functional analyses ^40,41,59,62^.

### Methodological contributions and validation

Our method builds on the premise that terminal branches (S.N. = 1) are preferential sites for synaptic connections, a hypothesis supported by prior studies^34,42–45^. By integrating this morphological feature with proximity thresholds derived from light microscopy resolution limits (∼5 *μ*m), we generated adjacency matrices that captured modular structures statistically aligned with EM-derived ground truth. The modest but significant correlations (Pearson: 0.05–0.215), Adjusted Rand Index (ARI: 0.04–0.6), and Normalized Mutual Information (NMI: 0.1–0.56) between inferred and EM-based connectomes underscore two key points: (1) spatial proximity and terminal branching hierarchy provide meaningful signals for approximating network architecture, and (2) these signals are insufficient to fully replicate synaptic-resolution connectivity. Importantly, the significant z-scores against randomized null models (p<0.05) confirm that inferred modules are non-random and retain biological relevance, even if incomplete.

### Biological insights and circuit mapping

In zebrafish, our method identified brain-wide modules that recapitulated known functional circuits, such as retinotopic projections in the tectum and vestibulo-ocular reflex pathways involving the cerebellum. These findings align with prior anatomical and functional studies, suggesting that mesoscale connectomes derived from light microscopy data can serve as a scaffold for hypothesis generation. For instance, the hierarchical decomposition of modules into submodules revealed spatially segregated neuronal populations, hinting at specialized functional roles. However, the distributed nature of many modules—spanning multiple brain regions—highlights the challenge of disentangling tightly interconnected circuits without synaptic-resolution data.

### Limitations and methodological considerations

While our approach offers scalability, several limitations must be acknowledged. First, the reliance on Peters’ rule (proximity as a proxy for connectivity) introduces both false positives and negatives, particularly in dense neuropil regions where axons and dendrites overlap without forming synapses. This is reflected in the modest validation metrics and underscores the need for complementary molecular or functional data to refine predictions. Second, the assumption that synaptic partners predominantly localize to SN = 1 branches may not generalize across species. For example, mammalian neurons often exhibit synaptic distributions across higher-order branches, a limitation highlighted by recent studies in mice. Third, the method’s performance is sensitive to registration accuracy and sampling density. In the zebrafish dataset, which covers ∼3% of total neurons, undersampling likely skewed hub identification and intermodule connectivity patterns.

### Our method with respect to other works

Our work aligns with emerging efforts to extract network architecture from partial or mesoscale anatomical datasets. While traditional connectomics prioritizes synaptic-resolution EM, our method complements these approaches by enabling rapid, large-scale analyses when dense reconstructions are impractical. Similar to Kanari et al ^67^, we preserve essential network features, but with the added advantage of incorporating spatial constraints that better maintain known anatomical relationships. However, as Udvary et al. ^68^ demonstrated, axo-dendritic overlap alone poorly predicts connectivity in mammalian cortex, suggesting that integrating additional morphological features (e.g., bouton density, branch complexity) or molecular markers (e.g., neurotransmitter identity) could enhance accuracy. Future work should explore hybrid approaches that combine our scalable framework with these complementary features.

### Future directions

To address these limitations, future iterations of this framework could incorporate multi-modal data, such as functional imaging or transcriptomic profiles, to constrain connectivity predictions. Additionally, refining proximity thresholds using species-specific synaptic distance distributions or dynamically adjusting thresholds based on neuropil density may improve biological fidelity. The integration of generative models or graph neural networks could further bridge the gap between mesoscale approximations and synaptic-resolution connectomes. Finally, expanding validation to include more organisms and brain regions will clarify the method’s generalizability.

In summary, our method provides a pragmatic tool for approximating connectomes in datasets lacking synaptic annotation. By balancing scalability with biological interpretability, it enables exploratory analyses of brain-wide network organization and generates testable hypotheses for targeted EM or functional studies. While not a replacement for synaptic-resolution connectomics, this approach fills a critical niche in the era of large-scale neuroanatomical datasets, offering a pathway to extract circuit-level insights from existing resources.

## CONCLUSIONS

Our approach provides a scalable framework for approximating connectomes when synaptic data are unavailable, but its limitations including reliance on spatial proximity and variable performance across brain regions should be weighed against the need for synaptic-resolution studies in targeted circuits. Moreover, the organization of the modules obtained from the zebrafish larva light microscopy brain dataset shows that the application of such top-down approach of community detection is able to capture relevant circuit motifs and may be applied to several datasets at both the circuit and whole-brain scale.

## Methods

### Dataset and data extraction

We used a whole-brain atlas of the zebrafish larva ^40^. The atlas comprises more than 3000 neurons, approximately 3% of the total neurons in the zebrafish larvae brain, covering almost the entire brain, anatomically reconstructed by random and sparse Green Fluorescent Protein (GFP) labelling using BGUG ^69^ or with other labelling techniques. With a typical resolution of 300 x 300 x 1000 nm (X, Y, Z), the size of the skeletonized dataset was approximately 60 MB. Skeletonized neurons have been downloaded in the .swc format ^70^ from the mapzebrain website (https://mapzebrain.org/home).

### Computing the Strahler ordering for the neuronal anatomy reconstructions

For a given neuronal skeleton *i* from the original dataset of 3163 reconstructions, we imported the .swc file and used the ‘networkx’ library ^71^ (https://networkx.org/) to convert the corresponding collection of ordinated endpoints to a tree-like directed graph representing its anatomical structure. The soma is considered as a root and the connected endpoints as children. In order to build an approximated connectome from this dataset, we first classified each skeleton endpoint within its branching hierarchy using Strahler ordering ^41^. In this representation, terminal endpoints, those without branching children, obtain a value one. For the other endpoints in the .swc file, the integer value of the Strahler progressively increases with the number of upstream branches along the anatomical structure to reach the maximum at the level of the soma. For the calculation of the putative synaptic contacts and hence for the calculation of the connectome, we considered only endpoints with unitary Strahler numbers i.e. SN = 1 (below referred to as endpoint unless otherwise specified), which we denoted as 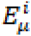 where the italic superscript refers to *i*-th neuron within the dataset and the Greek subscript stands for the *μ*-th endpoint of unitary Strahler number in the .swc file of *i*-th neuron.

### Leiden algorithm for neuronal module detection

We adopted the Leiden algorithm ^48^ to extract neuronal modules. The analysis of network parameters was performed in Python using the ‘leidenalg’ package (https://leidenalg.readthedocs.io/en/stable/index.html) .

The Leiden community detection algorithm iteratively refines an initial partition of nodes in a network to optimize a quality function called modularity Q. It has three main steps, which are fast local movement of nodes to communities, refinement of partition and lastly aggregation.

1. **Initialization:** The algorithm starts with a singleton partition which means each node is considered as its own community. The quality function Q is computed using the formulas below. We have chosen the well-known quality function for the undirected case^57^

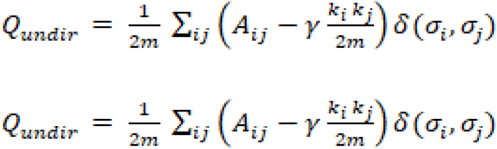

and for the directed case ^72^

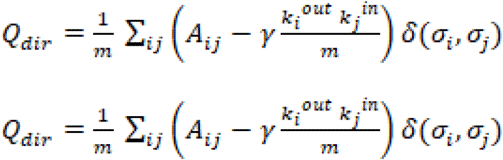

where *m* is total number of edges or total edge weight, *A* is the corresponding adjacency matrix for directed or undirected graph, γ >0 is the linear resolution parameter^59^ and is assumed equal to 1 in our analysis, higher the value of this parameter, larger the number of communities.*k*_*i* and *k*_*j* are the weighted node degree of node *i* and node *j* for the undirected case, while for the directed it is mentioned in superscript to specify whether it is weighted out-degree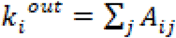 or weighted in-degree 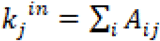, and finally *σ*_*i*_ denotes the community of node *i* with *δ* representing the Kronecker delta.
2. **Fast local movement of nodes:** A set of all nodes is created. The following steps are repeated until there are no further nodes in a list of nodes: (i) randomly select and remove a node from this set; (ii) for each community, calculate the gain in modularity as if the node were to move to that community; (iii) consider the move that results in the largest modularity gain, or no move if the modularity cannot be improved; (iv) update the partition by moving the node to the selected community; (v) all the neighboring nodes of the selected node which are not in the new community are added to the set of nodes. The steps are repeated until there is no node in the set of nodes.
3. **Refinement phase:** The partition P obtained from the fast local moving phase is refined as follows. Let the refined partition P* be initialized to a singleton partition, where each node represents a community. Randomly select one of the singleton communities in P*, and merge it to another community C* from P* with a probability that depends on the increase of the quality function. The higher the increase, the higher the probability of the merging. Repeat till all singleton community of P* are merged. Iterate these steps over all the communities of C belonging to P and obtain a new refined partition P*.
4. **Aggregation phase:** In this phase, each community in P* is considered as a node, while an edge between such communities is present if any of their nodes are connected. In this aggregated graph, communities are formed by merging all the aggregated nodes which are also subsets of the same community in the original partition.
5. **Convergence:** Return to step 2 of fast local moving of nodes and then terminate if there is no further change in the quality function.

### Comparison with the ground truth in Drosophila EM

We considered the EM annotated connectome of the Drosophila hemibrain as ground-truth and compared how close the reconstruction results when we applied on the same dataset our simplified method based on Strahler numbering and proximity range (SP method). For that, we identified eight modules using a community discovery approach on the EM connectome and considered the resulting partitions in submodules, 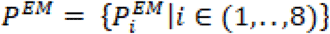 where 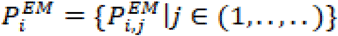 represents the *i*-module and 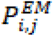 the *j*-sub-module associated with the module. Each 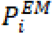 was considered for evaluating the sub-module partitions obtained based on the EM annotation 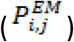 or retrieved with the approximated approach, 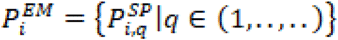, where 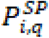 is the *q*-sub-module obtained with the SP method in association with the reference module 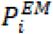 . For each module 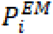, we calculated the corresponding *ARI*_*i*_ according to the following equation, as reported ^49,73^.

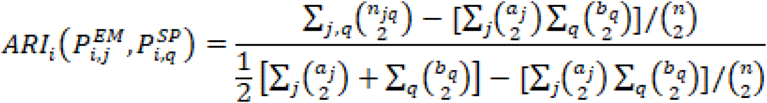

where *n*_*jq*_ represents the entries of the contingency matrix associated with the *i*-module, i.e. the number of neurons that are belonging to 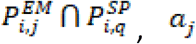, *a*_*j*_ and *b*_*q*_ are the sum of *n*_*jq*_ along *q* and *j* indices respectively, and *n* is the total number of partition elements. As for the NMI, we started by calculating the probability 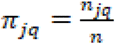, as the probability to belong simultaneously to 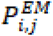 and,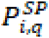 and the marginal distributions computed as 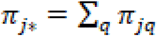 and 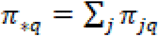 . We could then calculate the NMI as:

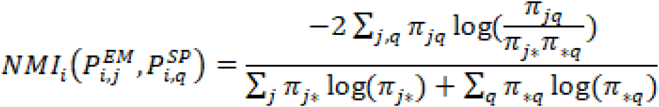

### Validation of ARI and NMI scores using Random null models

For validating the NMI and ARI parameters, we constructed 100 random networks for each of the mentioned 8 networks obtained with the SP method using two different approaches. In a first approach, called “degree-preserving shuffling or directed configuration”, we used directed_edge_swap function from the Python NetworkX library to swap the edges of the graph while keeping the node degree fixed. In a second approach, called Weight Randomization, we kept the original links between the nodes unchanged but changed the weight of the connection by randomly sampling from the original weight distribution. The z-score was calculated for ARI and NMI considering two different proximity range, 1 and 5 *μ*m, and the two types of random models as the difference of the between the actual value with respect to the average value obtained from the 100 null-model networks, divided by the corresponding standard deviation.

### Zebrafish undirected network reconstruction

We assume that all endpoints between two neurons form connections with each other. We introduce a proximity range *u* that represents the maximal effective distance at which two endpoints can form a connection. We set *u*=5*μm* based on the typical reconstruction error in optical microscopy ^40,74,75^. Then, a link between two neurons is present if at least two of their endpoints are at a distance smaller than *u*. That is, given two neurons *i,j* and their endpoints {*E*_*μ*_ ^(*i*)^ } and{*E*_*ν*_ ^(*j*)^ }, we compute the Euclidean distance matrix *d*_*μν*_ ^(*ij*)^= *d* (*E*_*μ*_ ^(*i*)^, *E*_*ν*_ ^(*j*)^), for all *μ*=1, ⋯,*N*_*i*_ and *ν*=1,⋯,*N*_*j*_. The elements of the resulting adjacency matrix *A*_*ij*_^*U*^ are the number of entries of *d*_*μν*_ ^(*ij*)^ such that *d*_*μν*_ ^(*ij*)^ ≤ *u* . Since *d*_*μν*_ ^(*ij*)^ is symmetric in the *ij* indexes, *A*_*ij*_^*U*^ is symmetric as well.

### Zebrafish directed network reconstruction

The morphological data we considered do not contain information between the dendritic and axonal endpoints. Therefore, given the endpoints {*E*_*μ*_ ^(*i*)^} of a neuron, we computed the radial distance *r*_*μ*_ ^(*i*)^ between each endpoint *E*_*μ*_ ^(*i*)^ and the soma. The associated probability distribution *Pi* (*r*) resulted often bimodal (Supplementary Figure 1), pointing to the fact that a large number of the endpoints are concentrated either close to the soma or far from it. We introduce a threshold *T* which approximately corresponds to the antimode of *P*_*i*_(*r*), i.e., the local minimum between the two modes. If 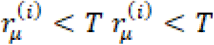, we assume that the corresponding synaptic endpoint is a dendritic endpoint for the neuron *i*, whereas 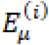 an axonal endpoint if 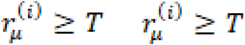.

Then, the connections from the *i*-th neuron to the *j* -th one are obtained from the distances 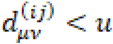, similarly to the undirected reconstruction. However, we now impose that a connection is counted only for those elements of 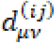 such that *μ* is an axonal endpoint for neuron *i*, and is a *ν* dendritic endpoint for neuron *j*. Clearly, the resulting adjacency matrix 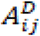 is not symmetric, since an axonal endpoint connecting to a dendritic endpoint does not imply that the reverse connection is present.

### Network properties

The analysis of network parameters ^76^ was performed using dedicated Python libraries. The following functions from the ‘networkx’ ^71^ (https://networkx.org/) library have been used: ‘connected_components’ to identify the network connected components, ‘average_shortest_path_length’ to compute the average shortest path length, ‘density’ to compute the density of the network, ‘average_clustering’ to compute the average clustering coefficient and ‘degree_assortativity_coefficient’ to compute assortativity correlation coefficient.

## Code and data availability

The scripts and data used to produce the results of the manuscript are available at https://github.com/dalMaschio-lab/Connectome_analysis and https://zenodo.org/records/15102704.

## Contributions

K.K.H.M. : conceptualization and data analysis.

M.B.: conceptualization, data analysis, anatomical analysis, supervision.

G.N. : conceptualization, data analysis, supervision.

S.S.: conceptualization and supervision.

M.D.M: conceptualization, supervision and funding acquisition.

All the authors participated in the writing, review and editing of the manuscript.

## Acknowledgments

We thank Delfina Iriarte, Physics department of University of Padua and Nora Nikoloska, Mathematics Department of University of Padua for their support in preliminary analyses; we thank Prof. Tomaso Erseghe from the Information Engineering Department of University of Padua for useful discussion. K.K.H.M. acknowledge support via a PHD scholarship from the Scuola Superiore Meridionale, Napoli, Italy.

## SUPPLEMENTARY FILE LEGENDS

**Supplementary Figure 1. Distribution of endpoints of neurons. A**) Typically, the distance between the endpoints of a neuron and the soma is distributed accordingly to two main distribution classes, following the Galtung classification ^77^. **B**) Bimodal class S. The antimode of the distribution is used to set a threshold (dashed line) to distinguish between dendrites and axon. C) Bimodal class U. The antimode of the distribution is used to set a threshold (dashed line) to distinguish between dendrites and axons.

**Supplementary Figure 2. The zebrafish module 1. A)** Anatomical representation of the high-order module U1. Colors are assigned according to the soma distribution in the brain regions. Scale bar = 100 *μ*m. **B)** Anatomical representation of the individual lower-order modules. Colors are assigned according to the location of the soma. Scale bar = 100 *μ*m.

**Supplementary Figure 3. The zebrafish module 2. A)** Anatomical representation of the high-order module U2. Colors are assigned according to the soma distribution in the brain regions. Scale bar = 100 *μ*m. **B)** Anatomical representation of the individual lower-order modules. Colors are assigned according to the location of the soma. Scale bar = 100 *μ*m.

**Supplementary Figure 4. The zebrafish module 4. A)** Anatomical representation of the high-order module U4. Colors are assigned according to the soma distribution in the brain regions. Scale bar = 100 *μ*m. **B)** Anatomical representation of the individual lower-order modules. Colors are assigned according to the location of the soma. Scale bar = 100 *μ*m.

**Supplementary Figure 5. The zebrafish module 5. A)** Anatomical representation of the high-order module U5. Colors are assigned according to the soma distribution in the brain regions. Scale bar = 100 *μ*m. **B)** Anatomical representation of the individual lower-order modules. Colors are assigned according to the location of the soma. Scale bar = 100 *μ*m.

**Supplementary Figure 6. The zebrafish module 6. A)** Anatomical representation of the high-order module U6. Colors are assigned according to the soma distribution in the brain regions. Scale bar = 100 *μ*m. **B)** Anatomical representation of the individual lower-order modules. Colors are assigned according to the location of the soma. Scale bar = 100 *μ*m.

**Supplementary Figure 7. The zebrafish module 7. A)** Anatomical representation of the high-order module U7. Colors are assigned according to the soma distribution in the brain regions. Scale bar = 100 *μ*m. **B)** Anatomical representation of the individual lower-order modules. Colors are assigned according to the location of the soma. Scale bar = 100 *μ*m.

**Supplementary Figure 8. Relationships between zebrafish modules**. Distribution of connections between the modules of the directed network. D: directed modules.

**Supplementary Figure 9. Proximity range and degree distribution comparisons**. Degree distribution plot for EM Drosophilia network and reconstructed network for Module 8 with proximity range defined **A)** 0.4 *μ*m **B)** 0.6 *μ*m. Measures to compare degree distributions for each proximity range varying between 0.2 *μ*m to 1.2 *μ*m **C)** KS Test **D)** JS divergence.

**Supplementary Table 1**. False positive and false negative analysis conducted between Drosophilia’s EM reconstructed module (ground truth) and its corresponding Strahler reconstructed module (predicted model) with proximity range fixed to 1*μ*m.

**Supplementary Table 2**. False positive and false negative analysis conducted between Drosophilia’s EM reconstructed **module 8** (ground truth) and its corresponding Strahler reconstructed **module 8** (predicted model) whose reconstruction parameter i.e. proximity range being varied between 0.2 *μ*m to 1.2 *μ*m.

**Supplementary Figure 10. The Drosophila module 1**. A) Anatomical reconstructions of neurons assigned to the high-order module one decomposed into the lower-order modules. Colors are assigned according to the order of the lower modules. Scale bar = 50 *μ*m.

**Supplementary Figure 11. The Drosophila module 2**. A) Anatomical reconstructions of neurons assigned to the high-order module two decomposed into lower-order modules. Colors are assigned according to the order of the lower modules. Scale bar = 50 *μ*m.

**Supplementary Figure 12. The Drosophila module 3**. A) Anatomical reconstructions of neurons assigned to the high-order module three decomposed into the lower-order modules. Colors are assigned according to the order of the lower modules. Scale bar = 50 *μ*m.

**Supplementary Figure 13. The Drosophila module 4**. A) Anatomical reconstructions of neurons assigned to the high-order module four decomposed into the lower-order modules. Colors are assigned according to the order of the lower modules. Scale bar = 50 *μ*m.

**Supplementary Figure 14. The Drosophila module 5**. A) Anatomical reconstructions of neurons assigned to the high-order module five decomposed into the lower-order modules. Colors are assigned according to the order of the lower modules. Scale bar = 50 *μ*m.

**Supplementary Figure 15. The Drosophila module 6**. A) Anatomical reconstructions of neurons assigned to the high-order module six decomposed into the lower-order modules. Colors are assigned according to the order of the lower modules. Scale bar = 50 *μ*m.

**Supplementary Figure 16. The Drosophila module 7**. A) Anatomical reconstructions of neurons assigned to the high-order module seven decomposed into the lower-order modules. Colors are assigned according to the order of the lower modules. Scale bar = 50 *μ*m.

**Supplementary Figure 17. The Drosophila module 8**. A) Anatomical reconstructions of neurons assigned to the high-order module seven decomposed into the lower-order modules. Colors are assigned according to the order of the lower modules. Scale bar = 50 *μ*m.

**Supplementary Figure 18**. Z score computed for Pearson correlation between EM module 6 and 8 and their corresponding Strahler reconstructed counterparts (shown in black markers). The blue markers are the z-scores computed between EM modules and the degree preserving randomized networks of those modules.

