## supplemental figures, tables and text for "A method for extracting an approximated connectome from libraries of single cell anatomical reconstructions": Supporting Information.docx

**Validation of ARI and NMI scores using Random null models, details**

We considered the 8 networks from the SP method and let’s represent them as
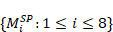
. Before proceeding to compute the z-scores, we constructed 100 random networks for each of the mentioned 8 networks of SP method using two unique random null models:

1. Degree-preserving shuffling or **directed configuration** ^75^**:** In this method, we use *directed_edge_swap()* function of NetworkX library of python to Swap three edges in a directed graph while keeping the node degrees fixed. This method considers randomly chosen three connected edges such that
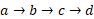
 and after the swap is applied it becomes
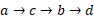
. In this way, we reach all possible states with the same in- and out-degree distribution in a directed graph and finally giving us directed configuration random null model.
2. **Weight randomization:** In this method, while keeping the links between the nodes unchanged or un-swapped, we assign weight to each edge by sampling from the weight distribution of all existing edge weights.

We computed the z-score for ARI and NMI for the 8 networks reconstructed with 1um proximity range and 5 μm proximity range using SP method. The mathematical formulation is as follows:

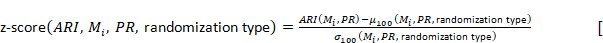
1]

where the notations refer to:

-
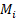
 - refers to the module(set of nodes belonging to community
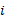

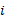
) for which we compute the similarity measures
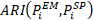
 or
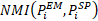
.
- PR - refers to proximity range which is either 1 μm or 5 μm used for reconstructing the network
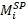
 using SP method.
- randomization type - refers to the type of random null model used to randomize the network
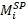
 i.e. directed configuration model or weight randomization.
-
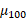
 and
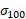
 are the mean and standard deviations over ARI or NMI computed on 100 random networks. Specifically, it is
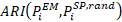
 or
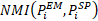

The above formula similarly can be written by replacing it with NMI measure.

#### **p-values computation**

We have computed the p-values for the z scores(Eq. [1]) obtained for ARI or NMI in two ways:

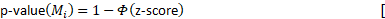
2]

- where
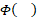
 refers to cumulative distribution function of standard normal distribution.

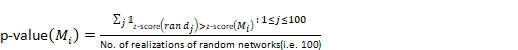
[3]

where
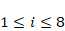
.

However, this approach diverges from our primary methodology, which selects parameters (proximity range = 1–5 µm, Strahler number = 1) based on biological and neuroscientific literature ^42–45^.

While degree distribution metrics like the KS test and JS divergence could theoretically guide parameter selection—particularly for out-degree distributions in directed networks—our approach prioritizes biologically grounded constraints. This preliminary analysis underscores the potential of distributional overlap measures for refining parameter choices in future work, though their applicability may depend on the specific network properties under study.
