## Supplementary figures and images for "A method for extracting an approximated connectome from libraries of single cell anatomical reconstructions"

### Supplementary Figure 1.png

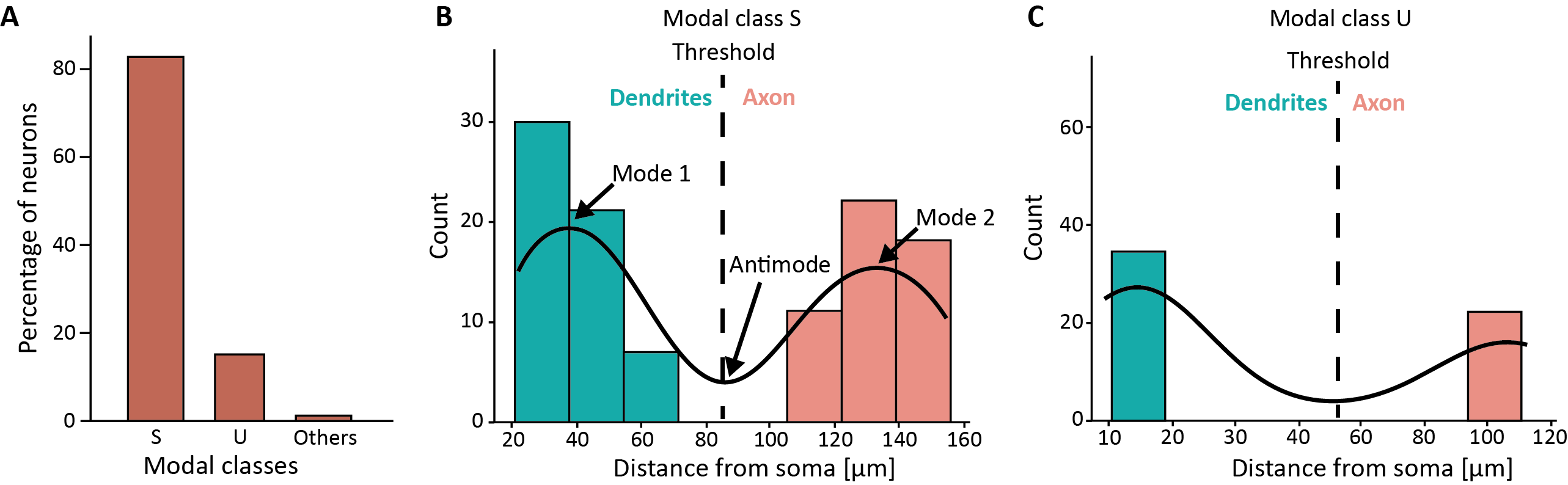

### Supplementary Figure 2.png

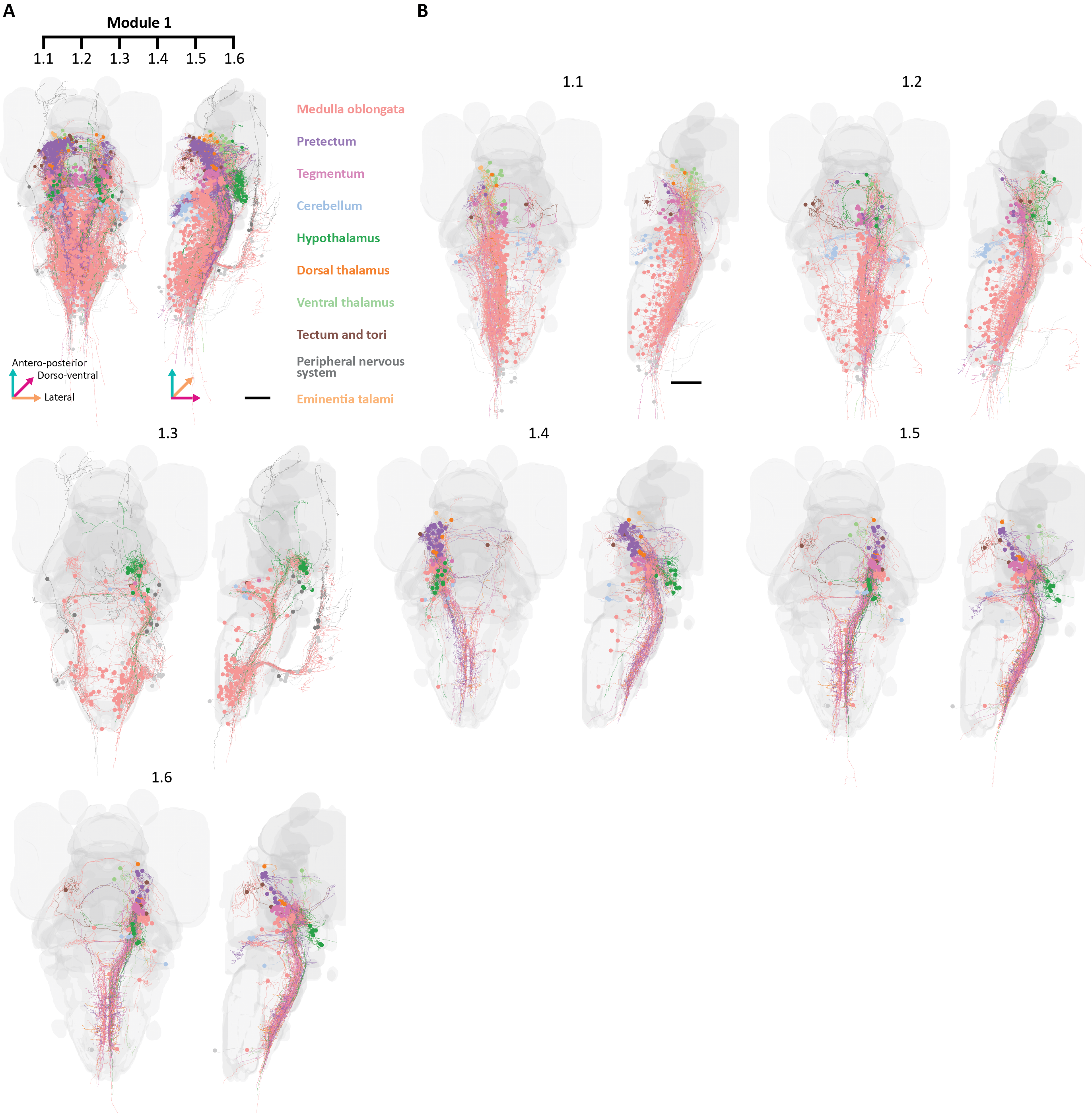

### Supplementary Figure 3.png

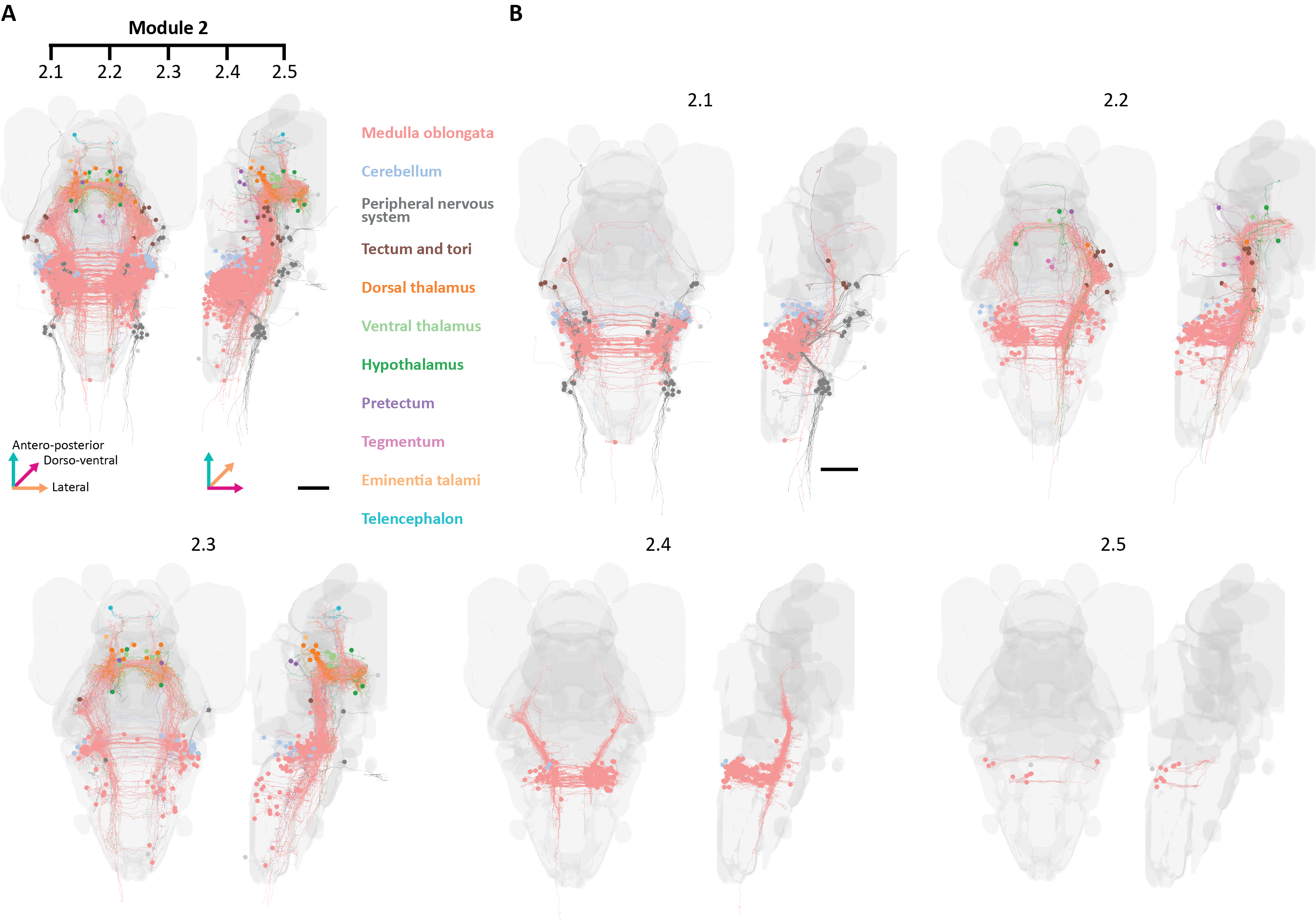

### Supplementary Figure 4.png

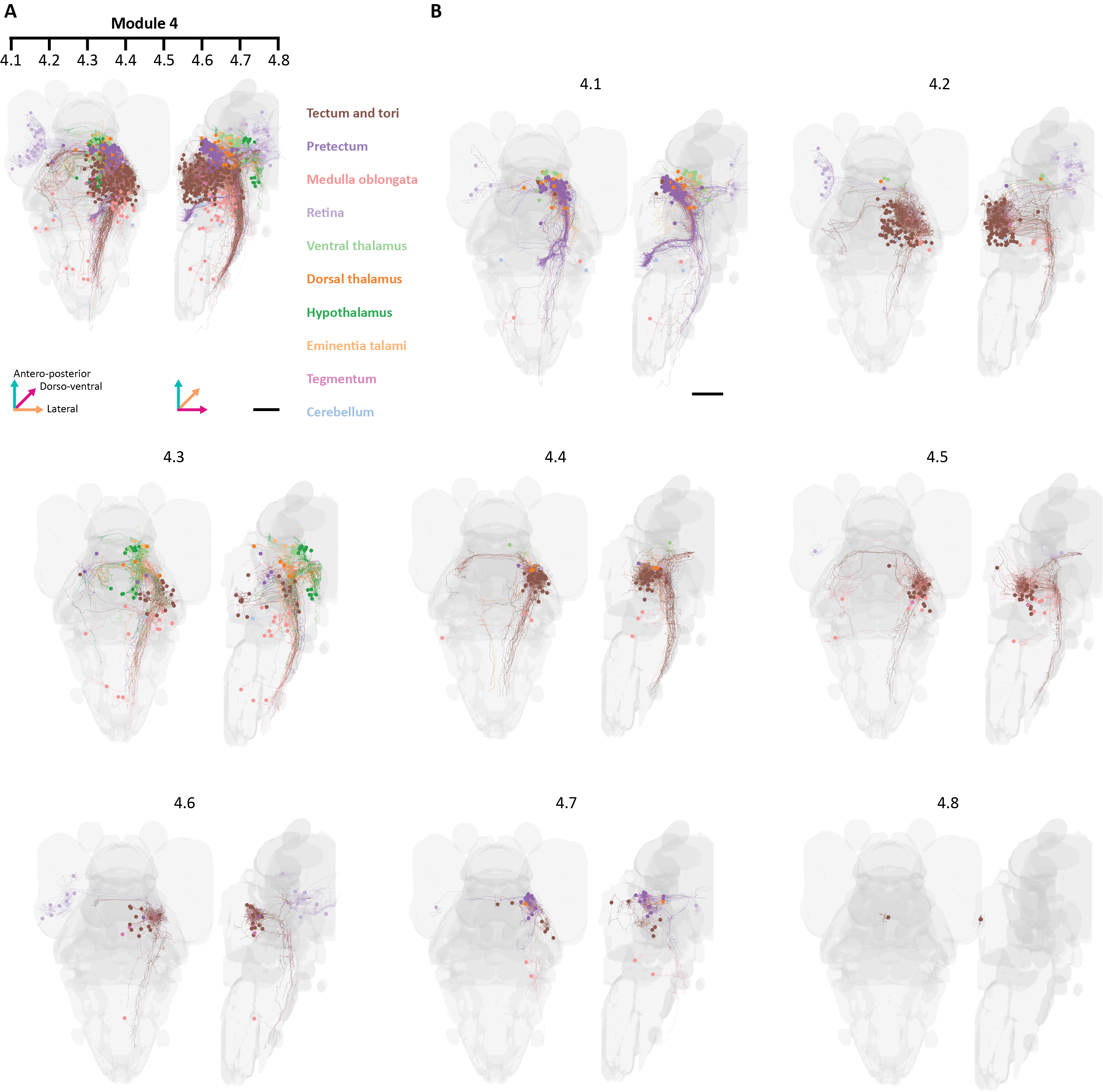

### Supplementary Figure 5.png

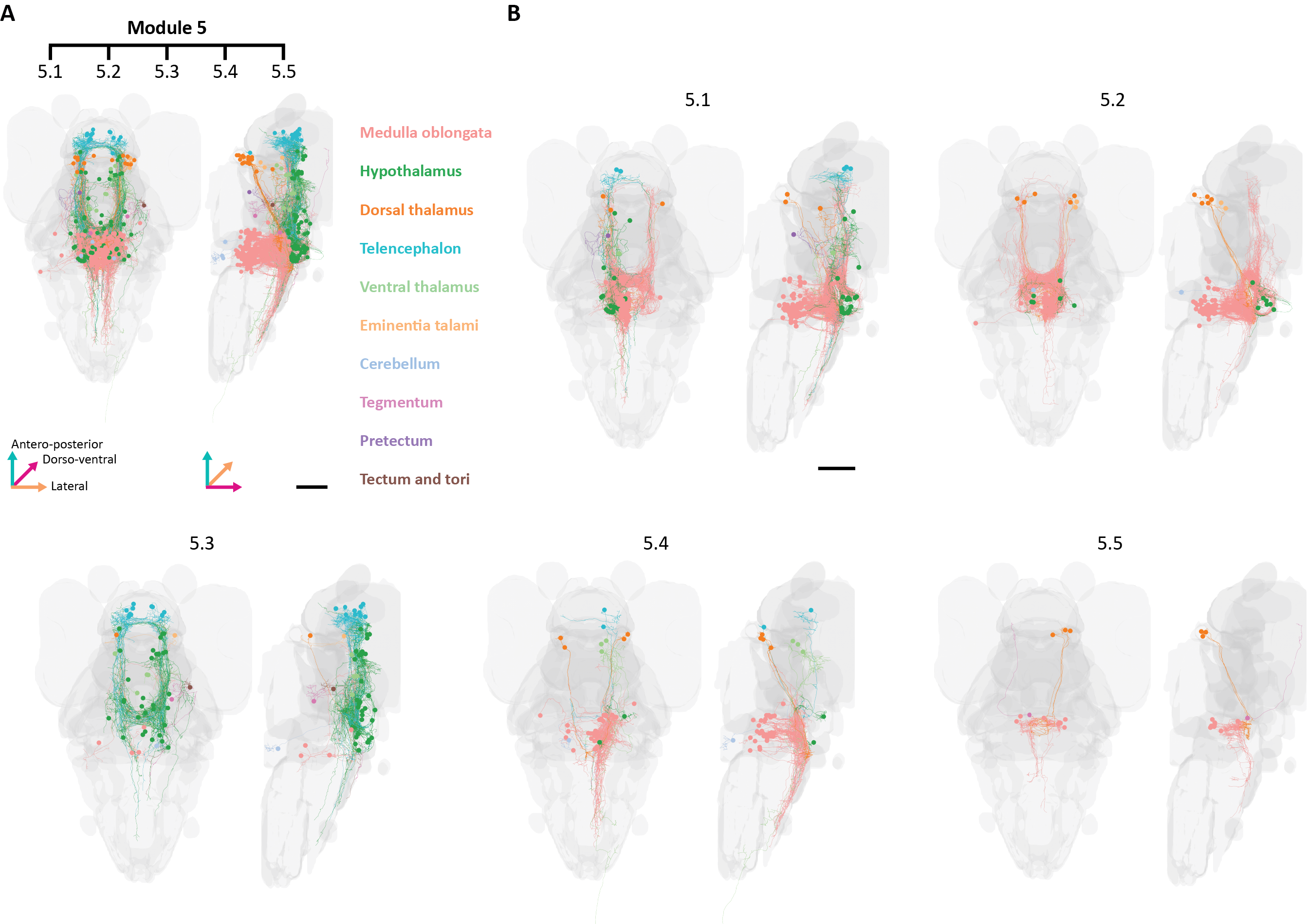

### Supplementary Figure 6.png

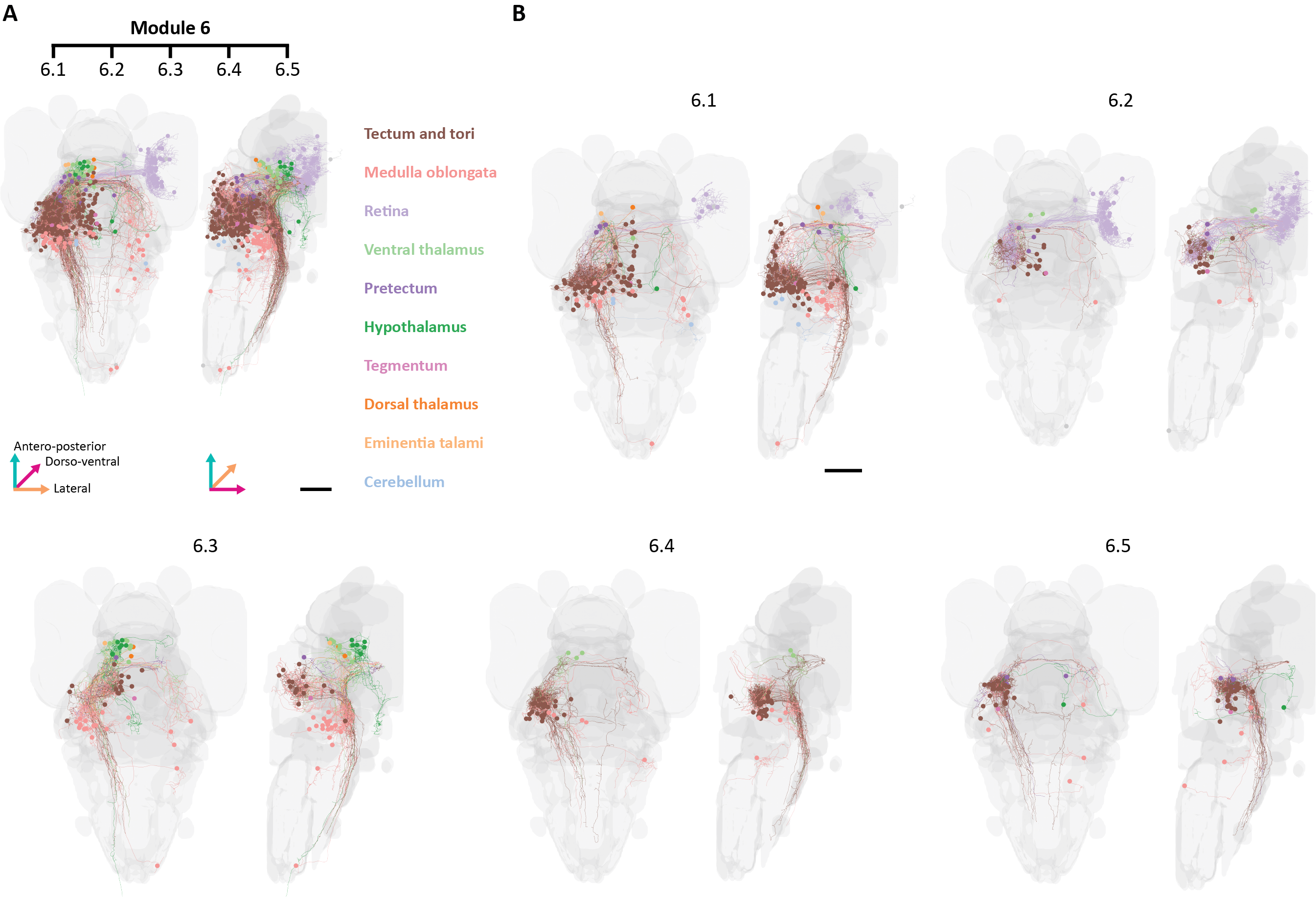

### Supplementary Figure 7.png

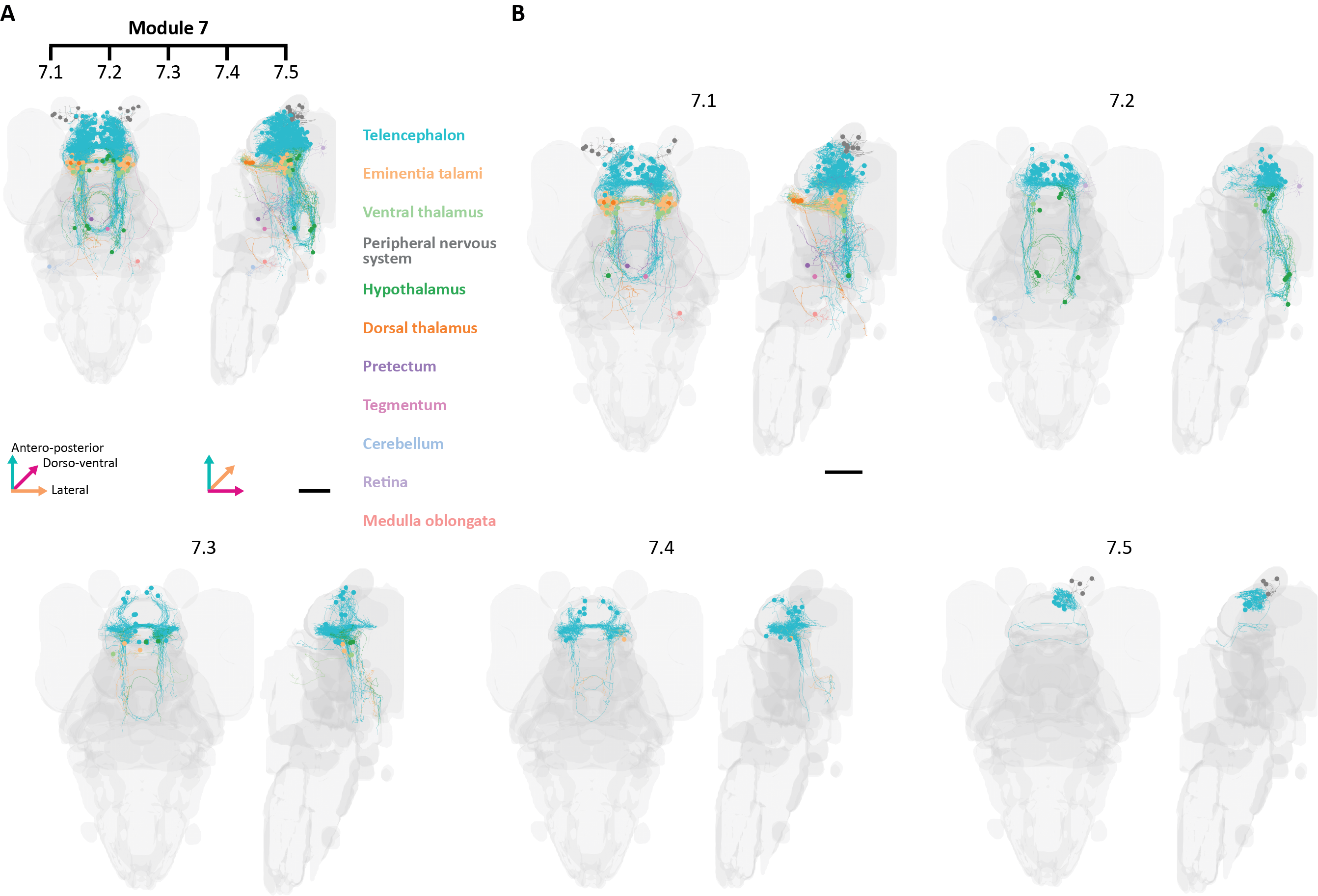

### Supplementary Figure 8.png

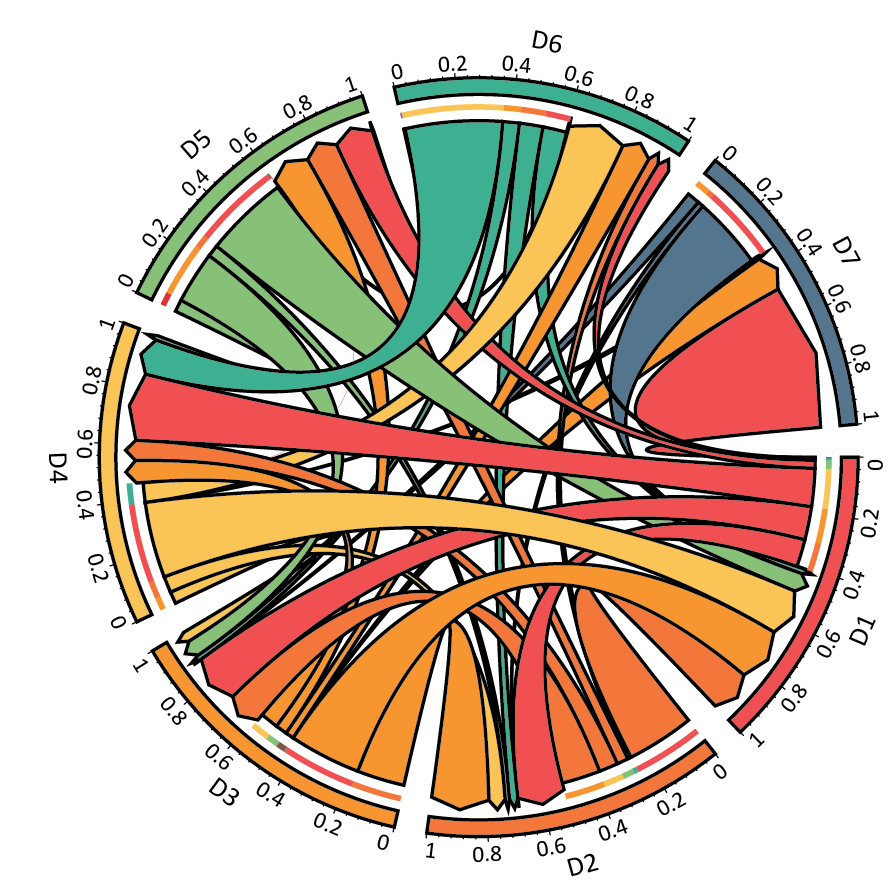

### Supplementary Figure 9.png

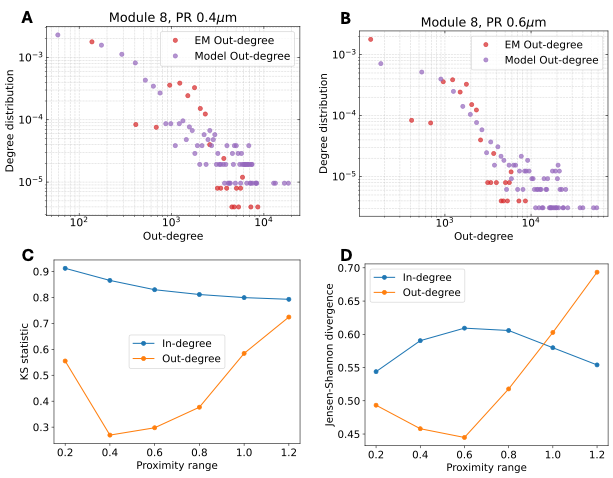

### Supplementary Figure 10.png

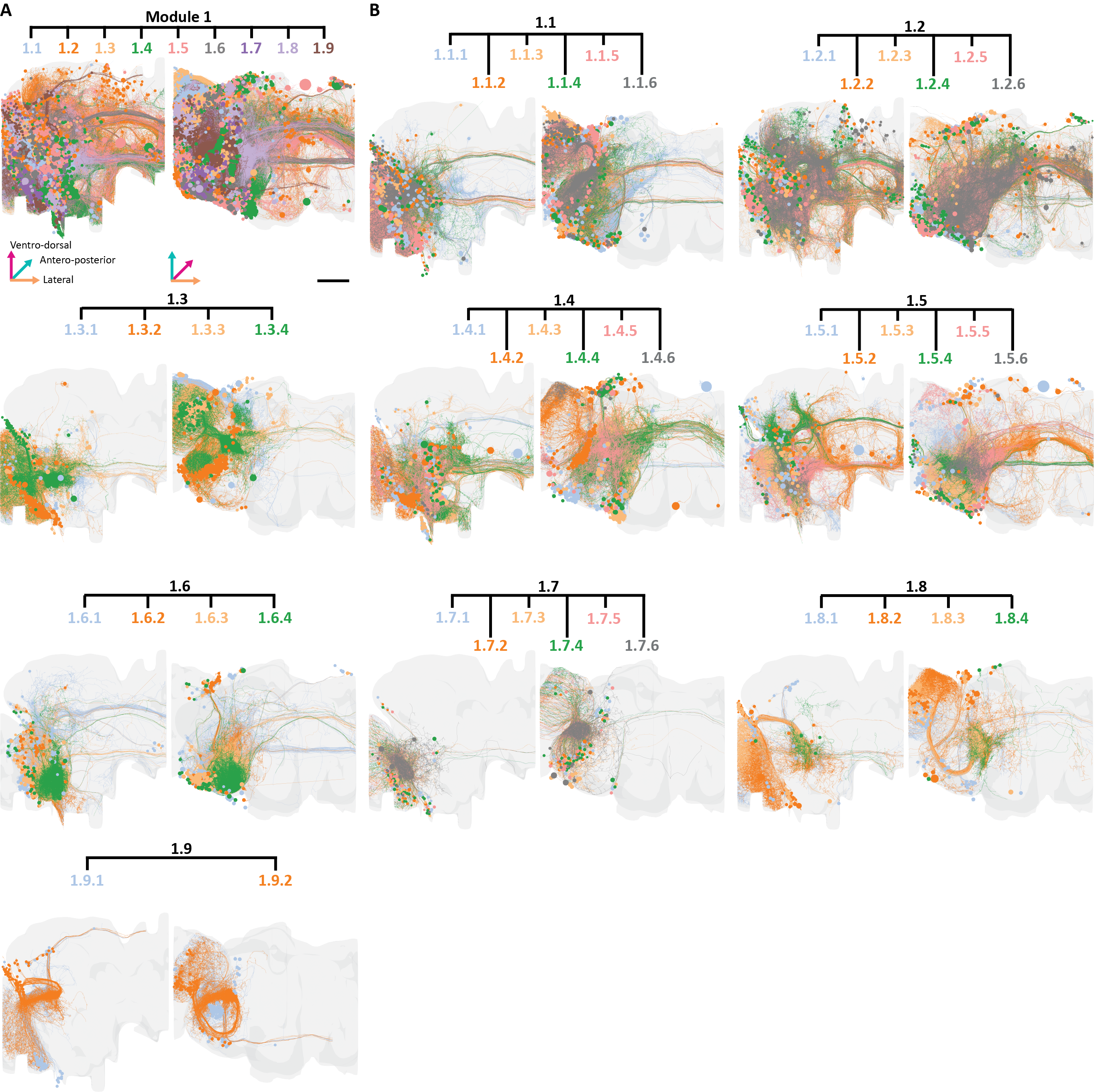

### Supplementary Figure 11.png

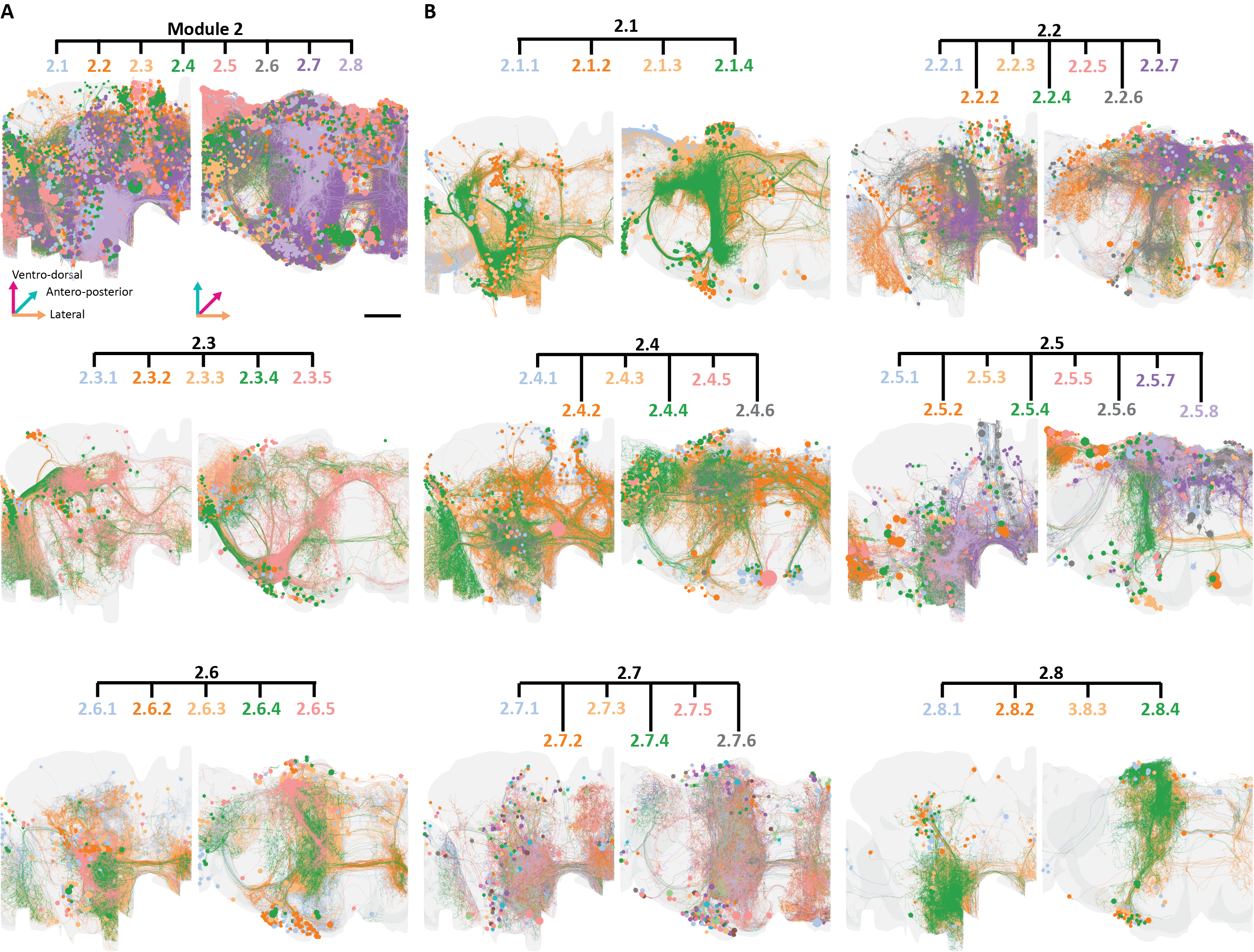

### Supplementary Figure 12.png

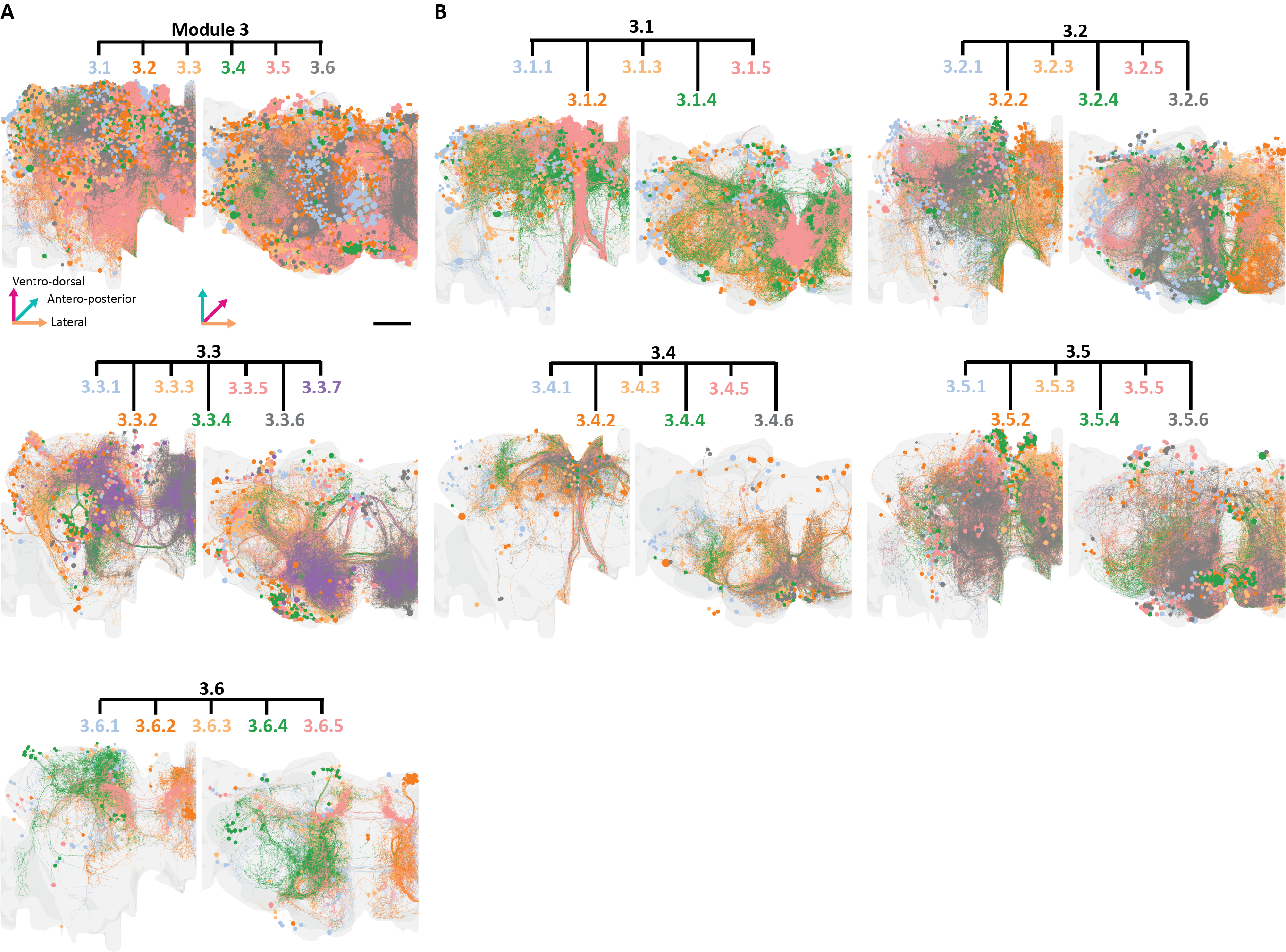

### Supplementary Figure 13.png

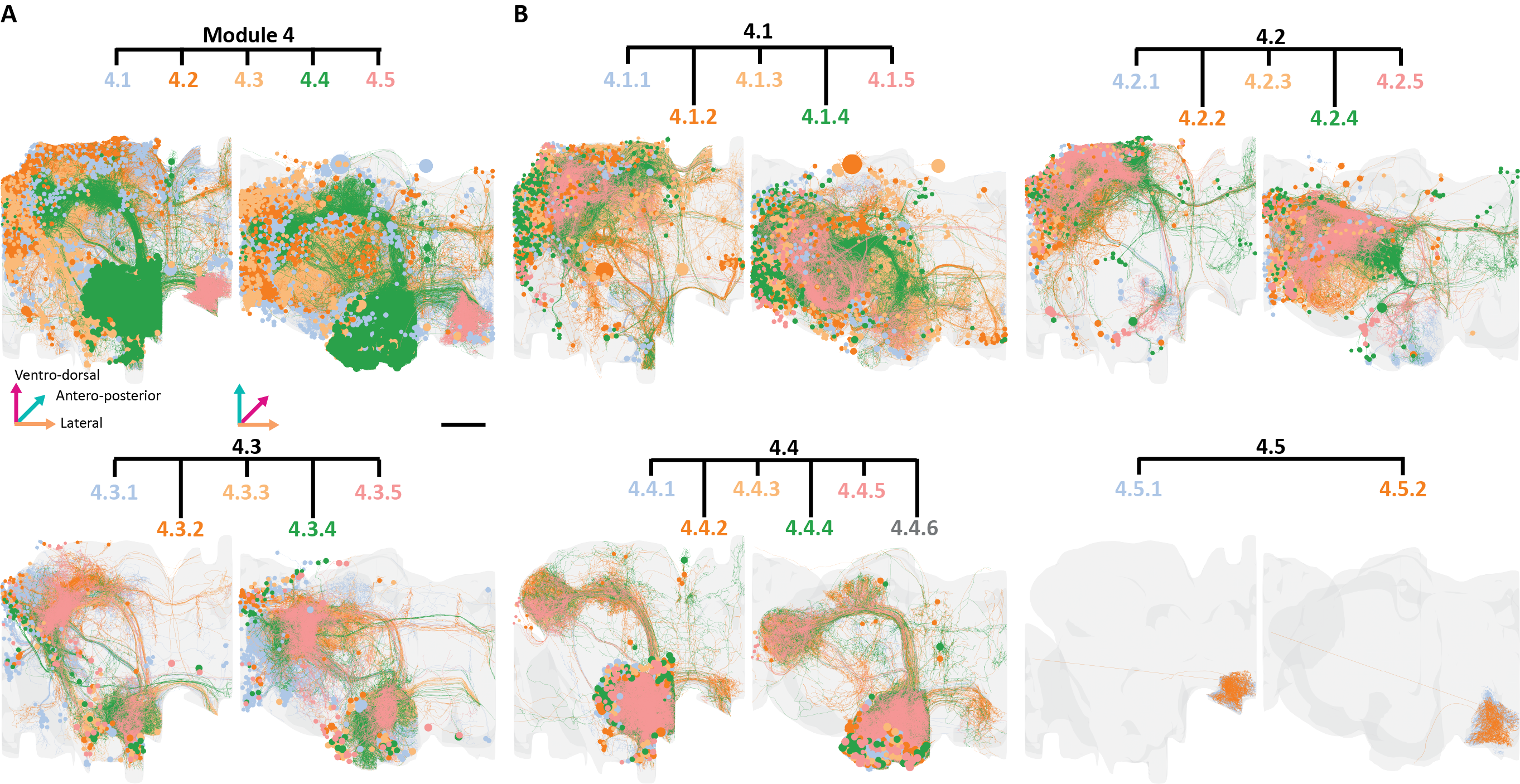

### Supplementary Figure 14.png

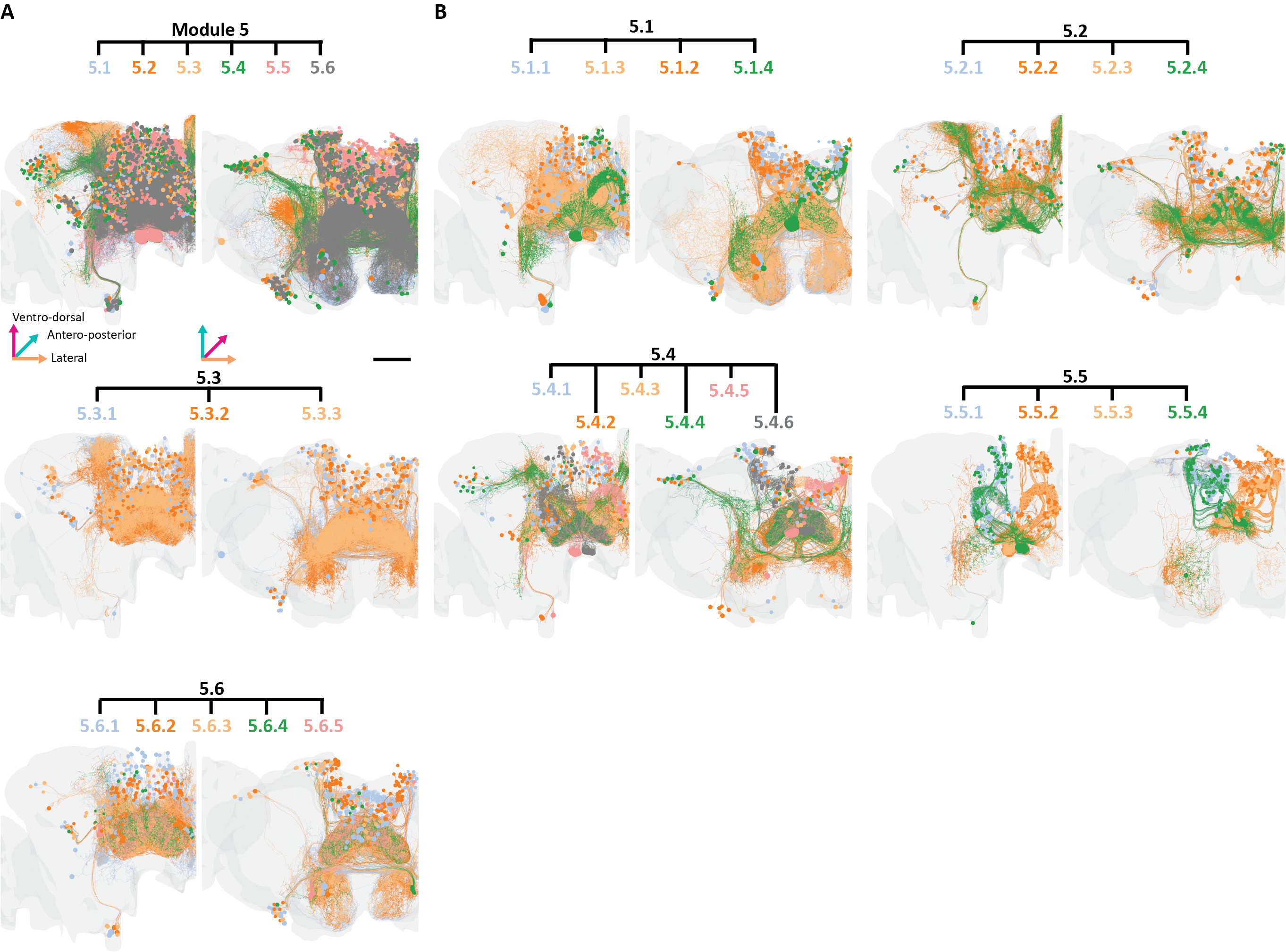

### Supplementary Figure 15.png

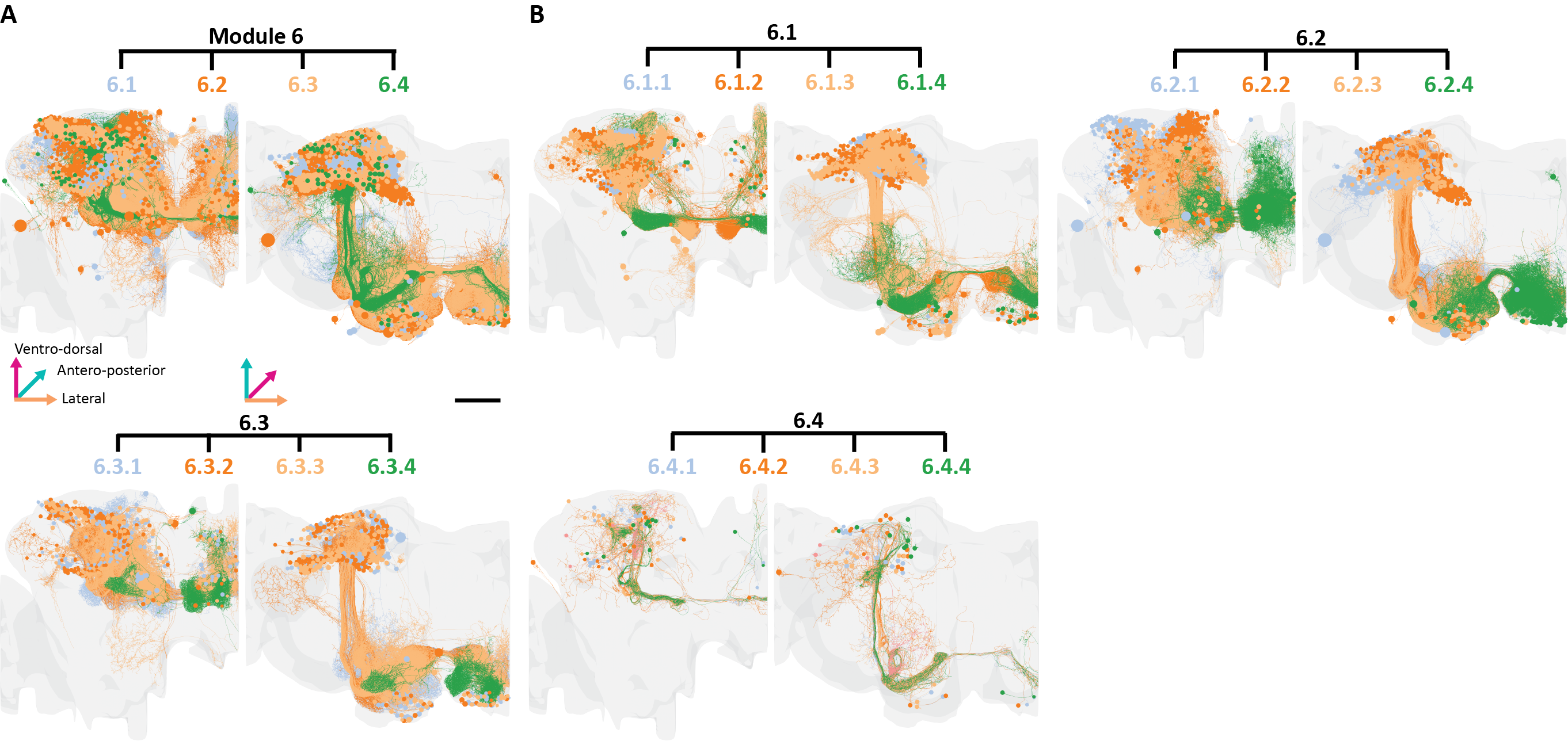

### Supplementary Figure 16.png

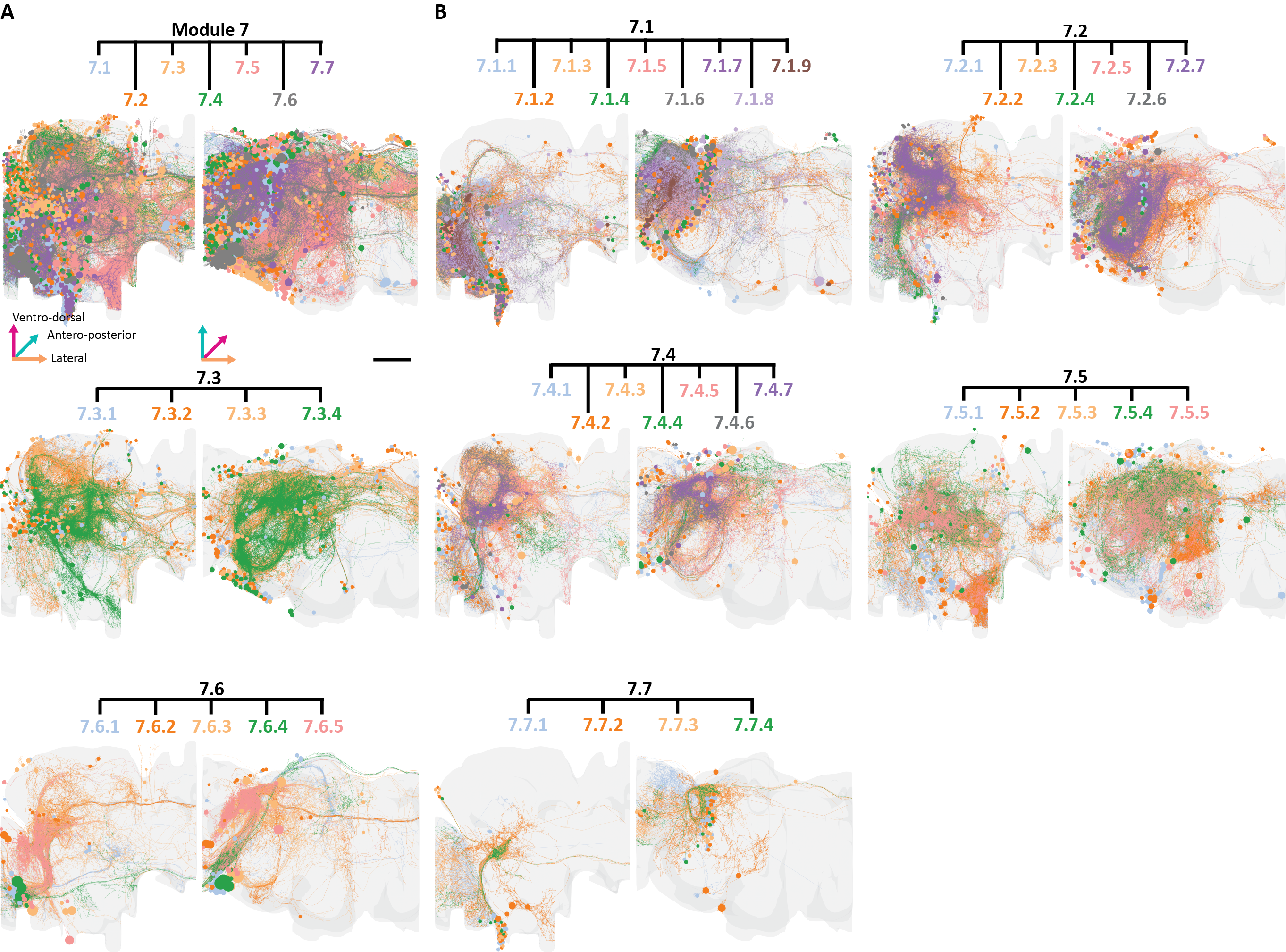

### Supplementary Figure 17.png

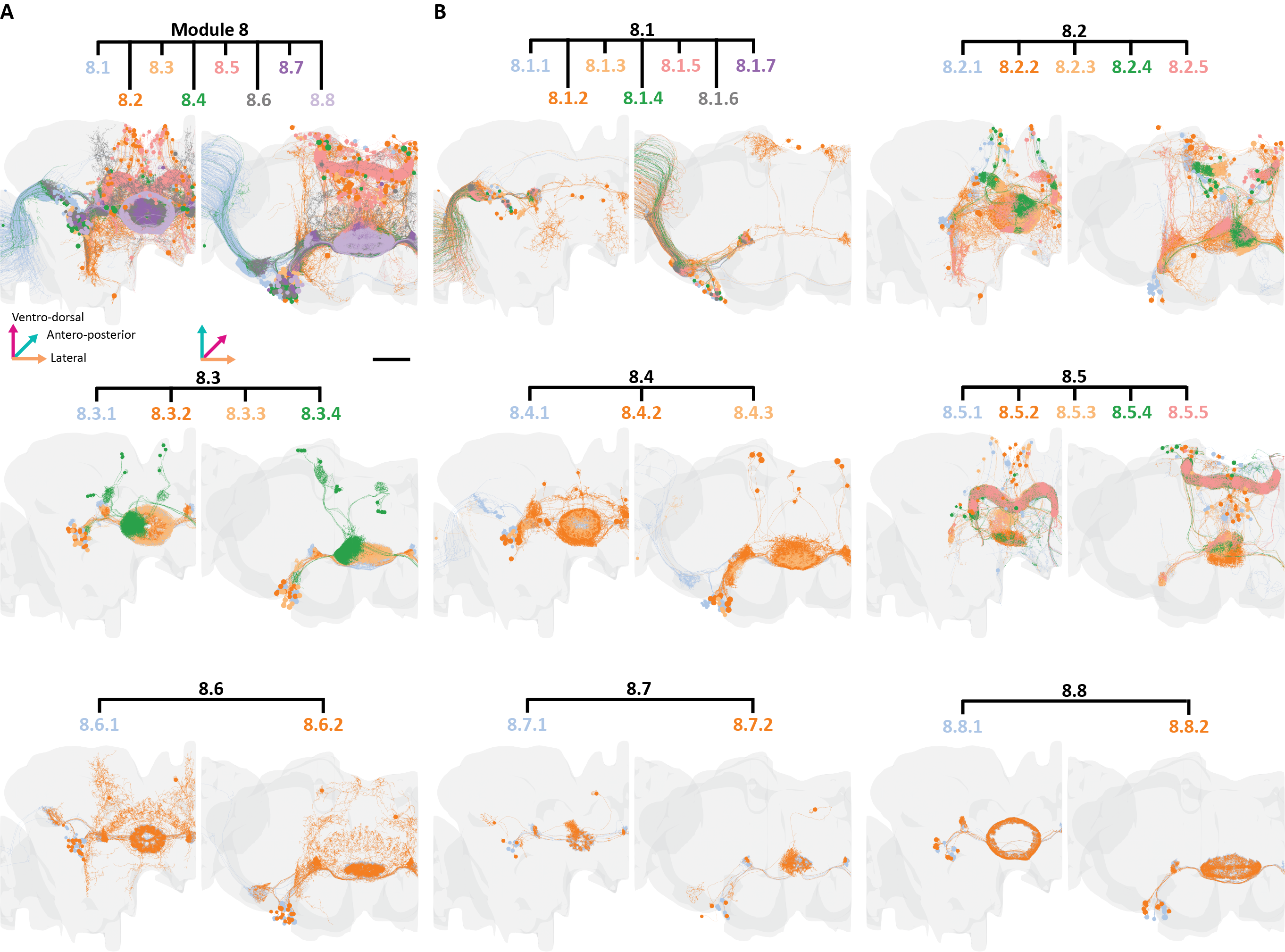

### Supplementary Figure 18.pdf

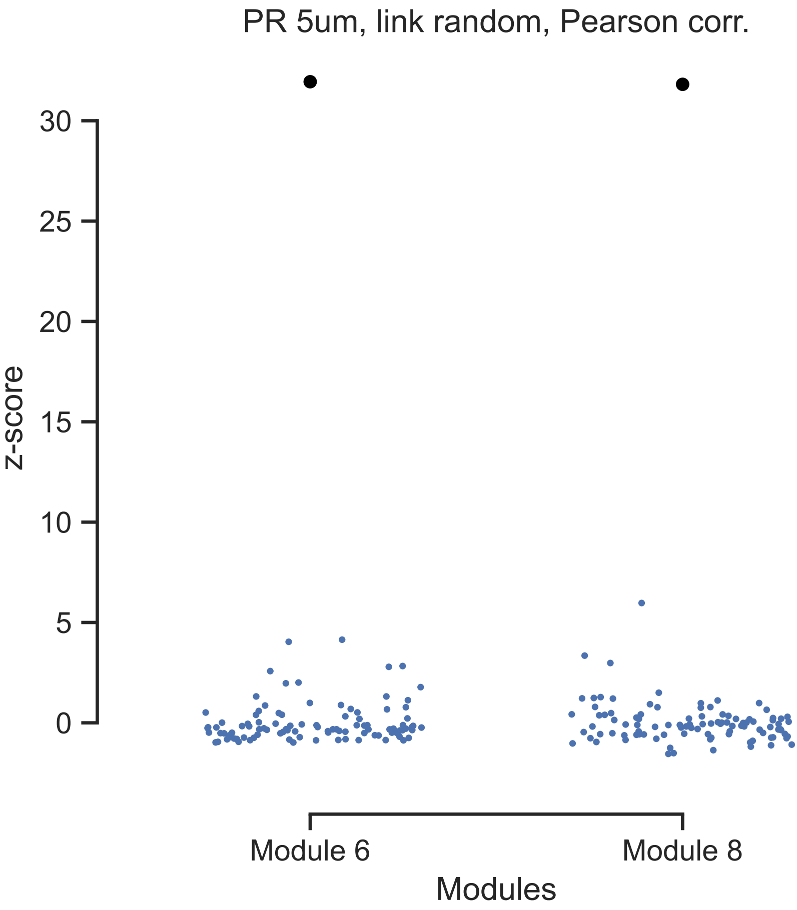

### Supplementary Table 1.tif

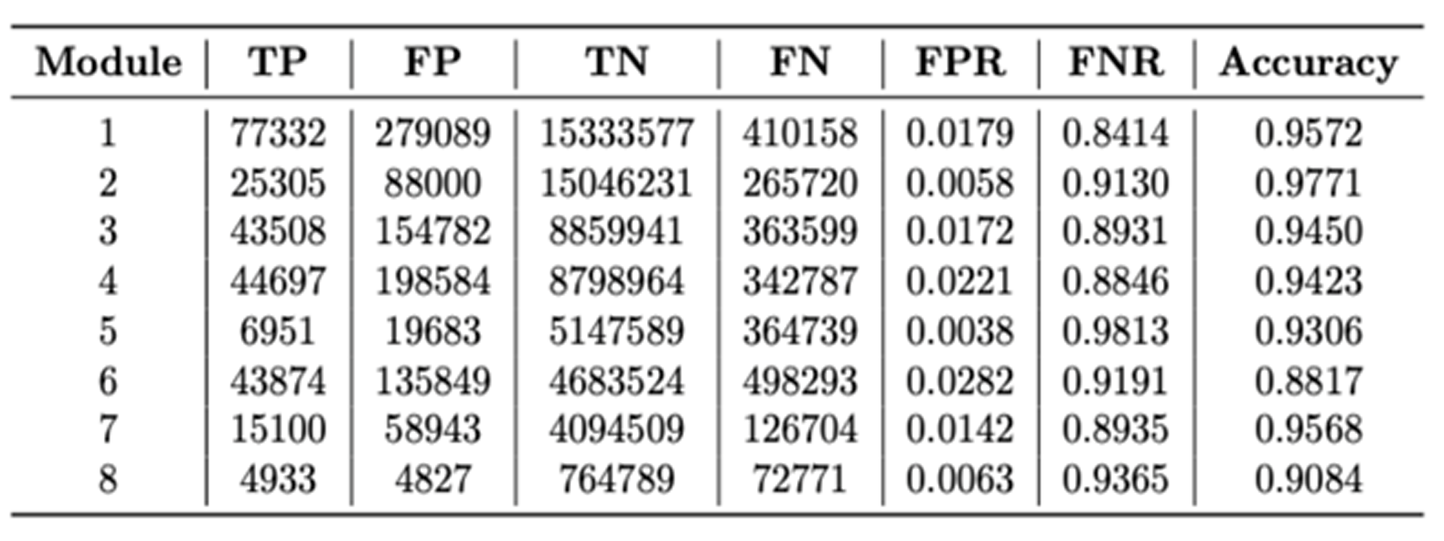

### Supplementary Table 2.tif

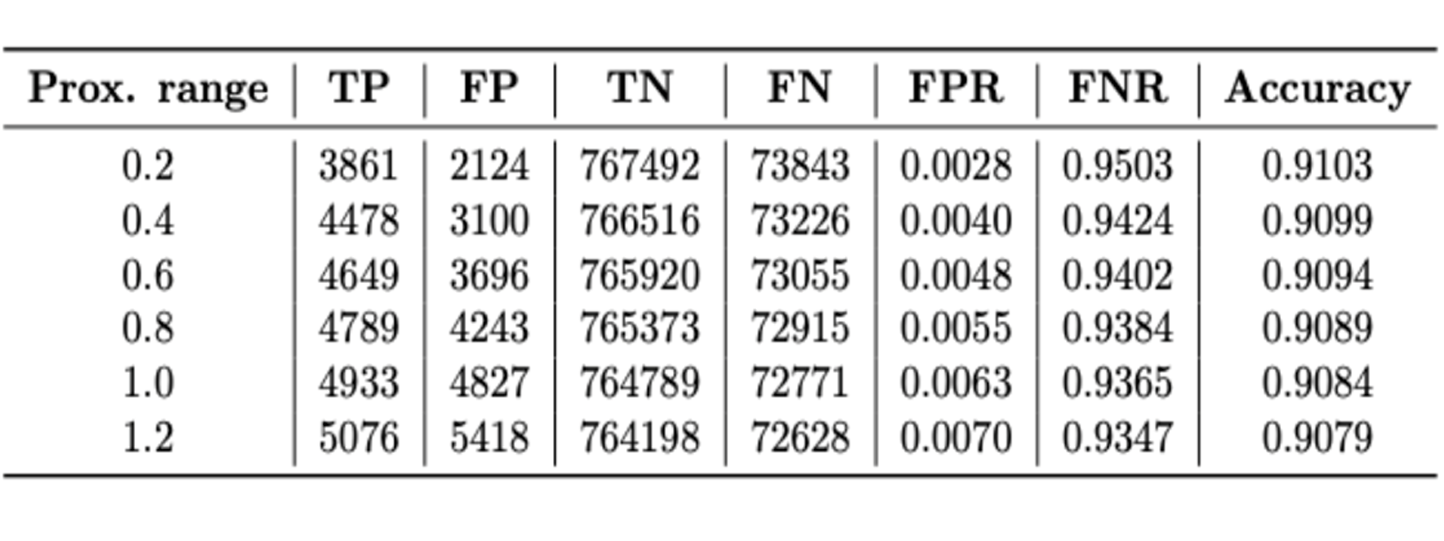
